# Universal activation mechanism of class A GPCRs

**DOI:** 10.1101/710673

**Authors:** Qingtong Zhou, Dehua Yang, Meng Wu, Yu Guo, Wangjing Guo, Li Zhong, Xiaoqing Cai, Antao Dai, Eugene Shakhnovich, Zhi-Jie Liu, Raymond C. Stevens, M. Madan Babu, Ming-Wei Wang, Suwen Zhao

## Abstract

Class A G protein-coupled receptors (GPCRs) influence virtually every aspect of human physiology. GPCR activation is an allosteric process that links agonist binding to G protein recruitment, with the hallmark outward movement of transmembrane helix 6 (TM6). However, what leads to TM6 movement and the key residue-level changes of this trigger remain less well understood. Here, by analyzing over 230 high-resolution structures of class A GPCRs, we discovered a modular, universal GPCR activation pathway that unites previous findings into a common activation mechanism, directly linking the bottom of ligand-binding pocket with G protein-coupling region. We suggest that the modular nature of the universal GPCR activation pathway allowed for the decoupling of the evolution of the ligand binding site, G protein binding region and the residues important for receptor activation. Such an architecture might have facilitated GPCRs to emerge as a highly successful family of proteins for signal transduction in nature.

## Introduction

GPCRs are membrane proteins that contain a seven-transmembrane helix (7TM) architecture^1–9^. In the last decade, we have witnessed a rapid development in GPCR structural biology (Figure 1a) and extensive research into the mechanism by which receptors are activated by diverse ligands including approved drugs^1–19^. While these studies have provided key insights into GPCR activation mechanism and implicated different parts of the receptor as being crucial for activation^10, 20–33^, they do not fully explain the pattern of conservation of residues and the number of disease-associated mutations that are known to map on distinct regions of the receptor (Figure 1—figure supplement 1). Although it is well established that outward movement of transmembrane helix 6 (TM6) upon ligand binding is a common feature of receptor activation^^3–5^, ^20–23^^, what leads to the movement of TM6, are they conserved, how the other helices are rearranged to facilitate this movement, and the key residue level changes of this trigger all remain less well understood (Figure 1b). Receptor activation requires global reorganization of residue contacts as well as water-mediated interactions^18–19^. Since prior studies primarily investigated conformational changes though visual inspection^20–22^ or through the presence or absence of non-covalent contacts between residues^8–10^, we reasoned that one could gain comprehensive knowledge about mechanism of receptor activation by developing approaches that can capture not just the presence or absence of a contact but also subtle, and potentially important alterations in conformations upon receptor activation.

**Figure 1.**
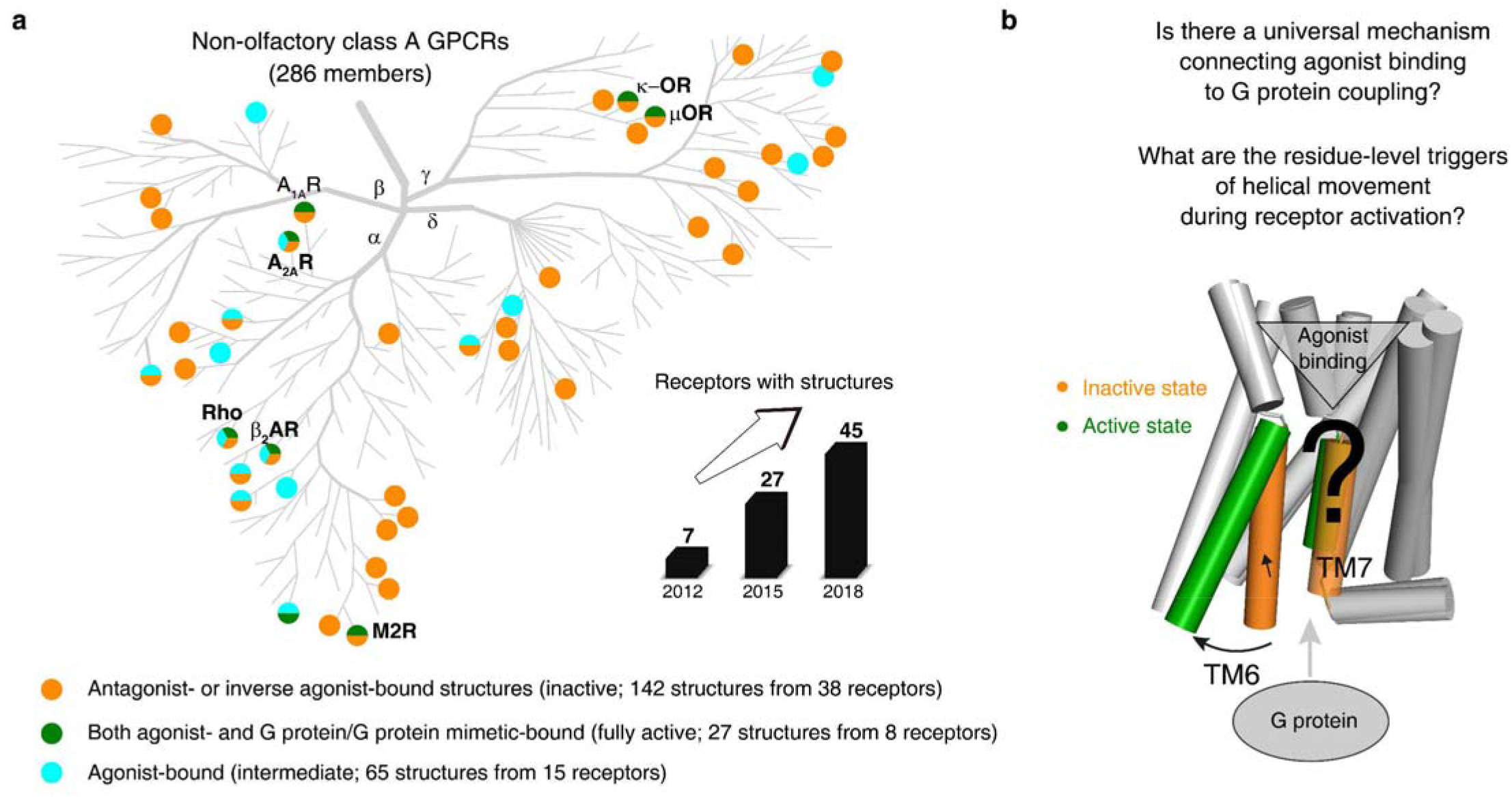
An increasing number of reported class A GPCR structures facilitates studies on universal activation mechanism. **a**, Distribution of structures in different states in the non-olfactory class A GPCR tree as of October 1, 2018. **b**, Universal GPCR activation mechanism and the residue-level triggers are not well understood. The following source data and figure supplement are available for figure 1: **Source data 1.** The released class A GPCR structures (as of October 1, 2018). **Source data 2.** Disease mutations occurred in class A GPCRs. **Figure supplement 1.** The pattern of conservation of residues and the map of number of disease-associated mutations on human class A GPCRs.

## Results

### A residue-residue contact score-based framework to characterize GPCR conformational changes

To address this, we developed an approach to rigorously quantify residue contacts in proteins structures and infer statistically significant conformational changes. We first defined a residue-residue contact score (RRCS) which is an atomic distance-based calculation that quantifies the strength of contact between residue pairs^34^ by summing up all possible inter-residue heavy atom pairs (Figure 2a and Figure 2—figure supplement 1a). We then defined ΔRRCS, which is the difference in RRCS of a residue pair between any two conformational states of a receptor that quantitatively describes the rearrangements of residue contacts (Figure 2b and Figure 2—figure supplement 1b). While RRCS can be 0 (no contact) or higher (stronger contact), ΔRRCS can be negative (loss in strength of residue contact), positive (gain in strength of residue contact) or 0 (no change in strength of residue contact). To capture the entirety of conformational changes in receptor structure upon activation, we computed the ΔRRCS between the active and inactive state of a receptor and defined two types of conformational changes (Figure 2c): (i) switching contacts: these are contacts that are present in the inactive state but lost in the active state (or *vice versa*) such as loss of intrahelical contacts between D/E^3×49^ (GPCRdb numbering^35^) and R^3×50^, and gain of interhelical hydrophobic contacts between residues at 3×40 and 6×48 upon receptor activation; and (ii) repacking contacts: these are contacts that result in an increase or decrease in residue packing such as the decreased packing of intrahelical sidechain contacts between W^6×48^ and F^6×44^, and the increase in interhelical residue packing due to the translocation of N^7×49^ towards D^2×50^ upon receptor activation. In this manner, we quantified the global, local, major and subtle conformational changes in a systematic way (*i.e.*, interhelical and intrahelical, switching and repacking contacts).

**Figure 2.**
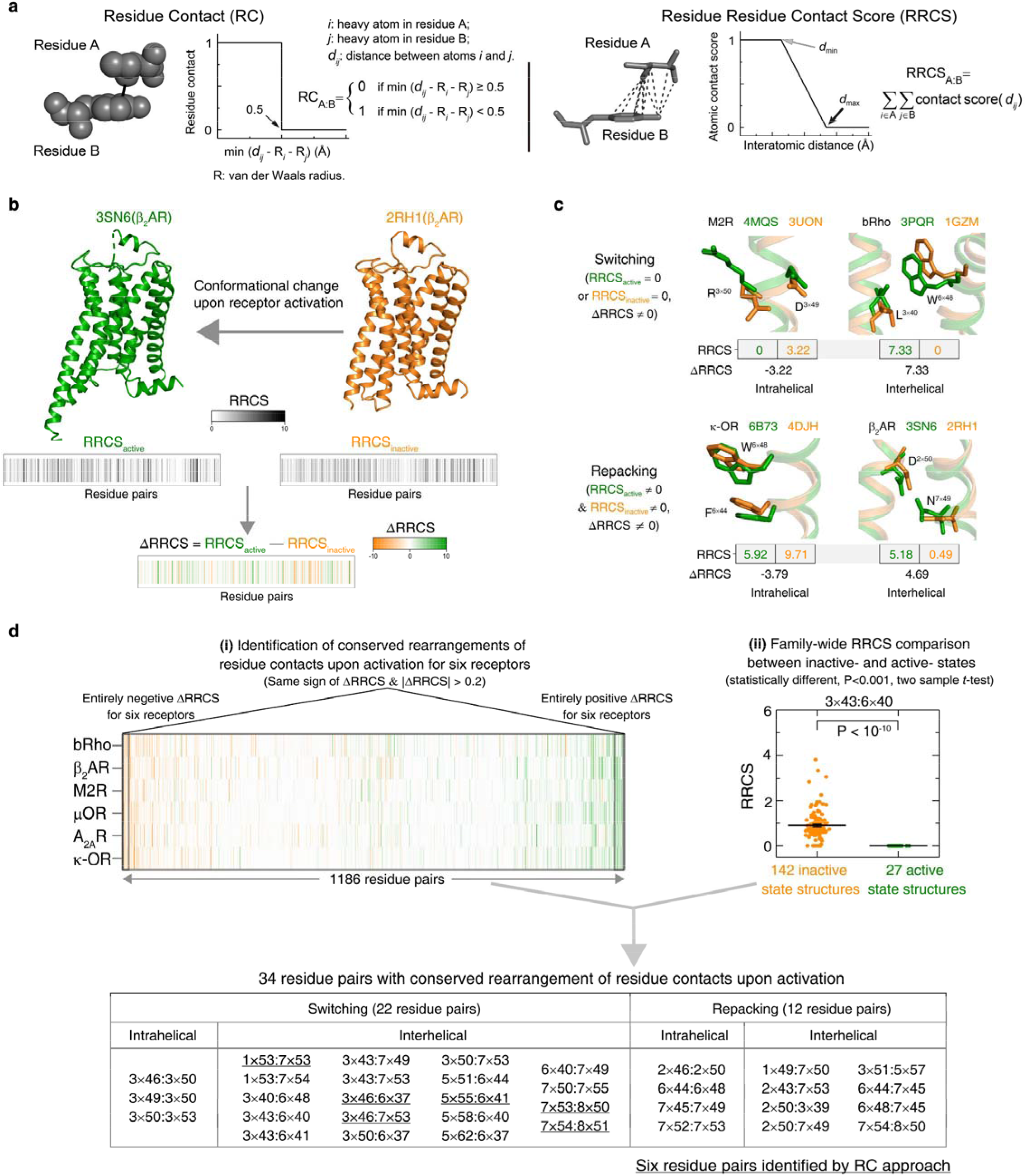
Understanding GPCR activation mechanism by RRCS and ΔRRCS. **a**, Comparison of residue contact (RC)^8^ and residue residue contact score (RRCS) calculations. RRCS can describe the strength of residue-residue contact quantitatively in a much more accurate manner than the Boolean descriptor RC. **b**, RRCS and ΔRRCS calculation for a pair of active and inactive structures can capture receptor conformational change upon activation. **c**, Two types of conformational changes (*i.e.*, switching and repacking contacts) can be defined by RRCS to quantify the global, local, major and subtle conformational changes in a systematic way. **d**, Two criteria of identifying conserved residue rearrangements upon receptor activation by RRCS and ΔRRCS. 34 residues pairs were identified based on the criteria (please see Methods, Figure 2—Source data 1 and 2 for details), only 6 of them were discovered before^8^. The following source data and figure supplement are available for figure 2: **Source data 1.** Calculated RRCS of 34 residue pairs constituting the universal activation pathway for released class A GPCR structures. **Source data 2.** Thirty-four residue pairs shown conserved rearrangements of residue contacts upon activation. **Figure supplement 1.** Calculation of RRCS and ΔRRCS.

We then analysed 234 structures of 45 class A GPCRs that were grouped into three categories (Figure 1a): (i) antagonist- or inverse agonist-bound (inactive; 142 structures from 38 receptors); (ii) both agonist- and G protein/G protein mimetic-bound (fully active; 27 structures from 8 receptors); and (iii) agonist-bound (intermediate; 65 structures from 15 receptors). Among them, six receptors [rhodopsin (bRho), β_2_-adrenergic receptor (β_2_AR), M2 muscarinic receptor (M2R), μ-opioid receptor (μOR), adenosine A_2A_ receptor (A_2A_R) and κ-opioid receptor (κ-OR)] have both inactive- and active-state crystal structures available. Given that ΔRRCS can capture major and subtle conformational changes, we computed RRCS for all structures and ΔRRCS for the six pairs of receptors and investigated the existence of a common activation pathway (*i.e.*, a common set of residue contact changes) across class A GPCRs. Two criteria (Figure 2d; further details in Methods) were applied to identify conserved rearrangements of residue contacts: (i) equivalent residue pairs show a similar and substantial change in RRCS between the active and inactive state structures of each of the six receptors (*i.e*., the same sign of ΔRRCS and |ΔRRCS| > cut-off for all receptors) and (ii) family-wide comparison of the RRCS for the 142 inactive and 27 active state structures shows a statistically significant difference (P<0.001; two sample *t*-test). This allowed us to reliably capture both the major rearrangements as well as subtle but conserved conformational changes at the level of individual residues in diverse GPCRs in a statistically robust and significant manner. Consistent with this, a comparison with earlier studies revealed that the RRCS based approach is able to capture a larger number of conserved large-scale and subtle changes in residues contacts (Figure 2d) that would have been missed by visual inspection or residue contact presence/absence criteria alone (see Methods for conceptual advance of this approach and detailed comparison)^8, 10, 20–22^.

### Discovery of the universal and conserved receptor activation pathway

Remarkably, for the first time, our analysis of the structures allowed the discovery of a universal and conserved activation pathway that directly links ligand-binding pocket and G protein-coupling regions in class A GPCRs (Figure 3). The pathway is comprised of 34 residue pairs (formed by 35 residues) with conserved rearrangement of residue contacts upon activation (Figure 2d), connecting several well-known but structurally and spatially disconnected motifs (CWxP^11, 20, 33^, PIF^3, 36^, Na^+^ pocket^19, 24, 33^, NPxxY^20, 23^ and DRY^11, 14, 37^) all the way from the extracellular side (where the ligand binds) to the intracellular side (where the G protein binds). Inspection of the rewired contacts as a ΔRRCS network reveals that the conserved receptor activation pathway is modular and involves conformational changes in four layers. In layer 1, there is a conserved signal initiation step involving changes in residue contacts at the bottom of the ligand-binding pocket and Na^+^ pocket. In layer 2, critical hydrophobic contacts are broken (*i.e.*, opening of the hydrophobic lock). In layer 3, microswitch residues (6×37, Y^7×53^) are rewired and in layer 4, the residue R^3×50^ and G protein contacting positions are rewired, making them competent to bind to G protein on the cytosolic side (Figure 3). Strikingly, recently released cryo-EM structures of three receptors (5-HT_1B_, A_1_R and μOR) in complex with G_i/o_ ^38^ also support the conservation of contacts involving these 34 residue pairs (Figure 4, Figure 4—figure supplements 1 and 2). These observations highlight the conserved and universal nature of a previously undescribed activation pathway linking ligand binding to G protein coupling, regardless of the subtypes of intracellular effectors (*i.e.*, G_s_, G_i/o_, arrestin or G protein mimetic nanobody/peptide, Figure 4a). Collectively, these findings illustrate how a combination of intrahelical and interhelical switching contacts as well as repacking contacts underlies the universal activation mechanism of GPCRs.

**Figure 3.**
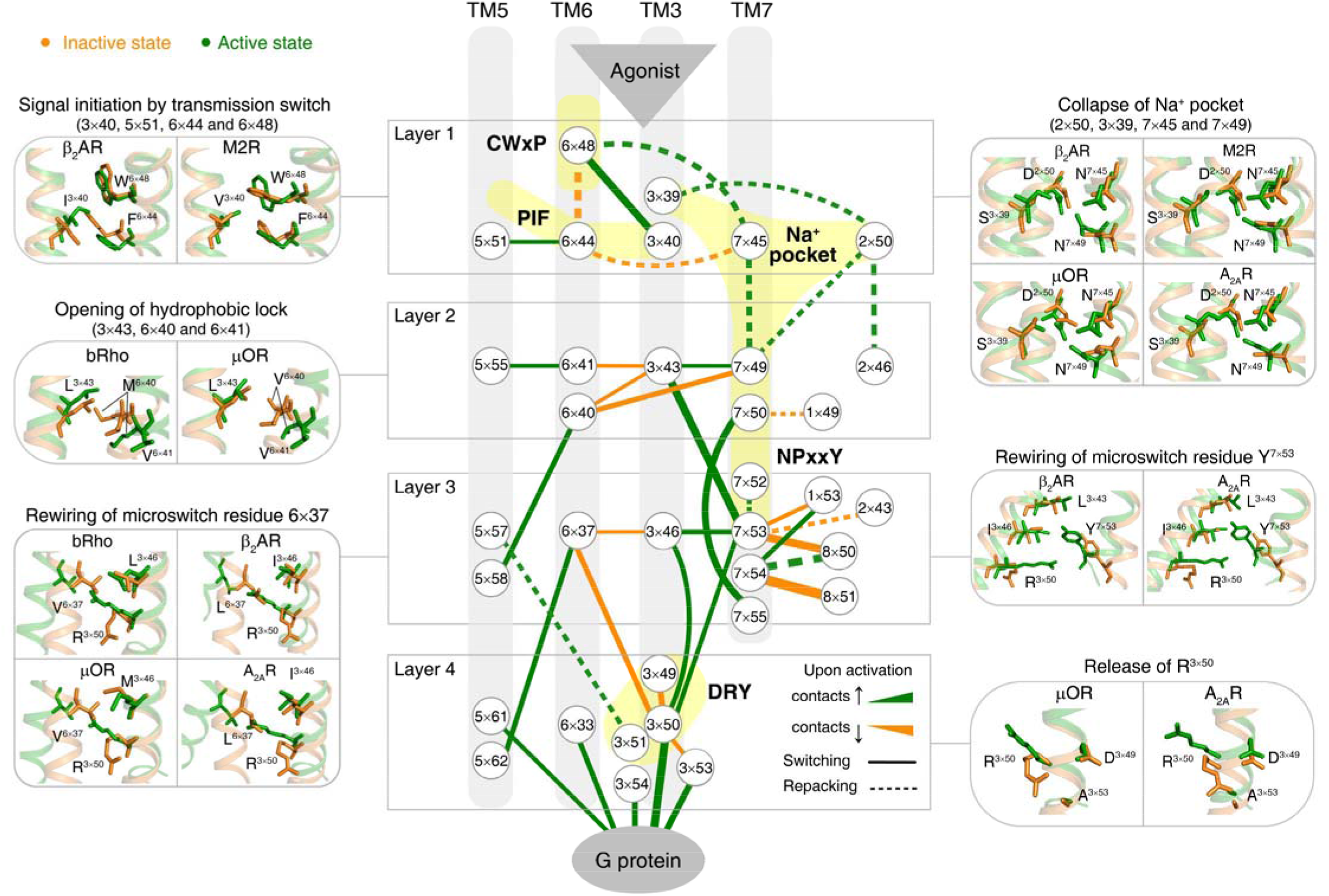
Universal activation pathway of class A GPCRs. Node represents structurally equivalent residue with the GPCRdb numbering^35^ while the width of edge is proportional to the average ΔRRCS among six receptors (bRho, β_2_AR, M2R, μOR, A_2A_R and κ-OR). Four layers were qualitatively defined based on the topology of the pathway and their roles in activation: signal initiation (layer 1), signal propagation (layer 2), microswitches rewiring (layer 3) and G protein coupling (layer 4). The following figure supplement is available for figure 3: **Figure supplement 1.** Rearrangements of ligand-residue contacts in ligand-binding pocket are not conserved, reflecting diverse ligand recognition modes.

**Figure 4.**
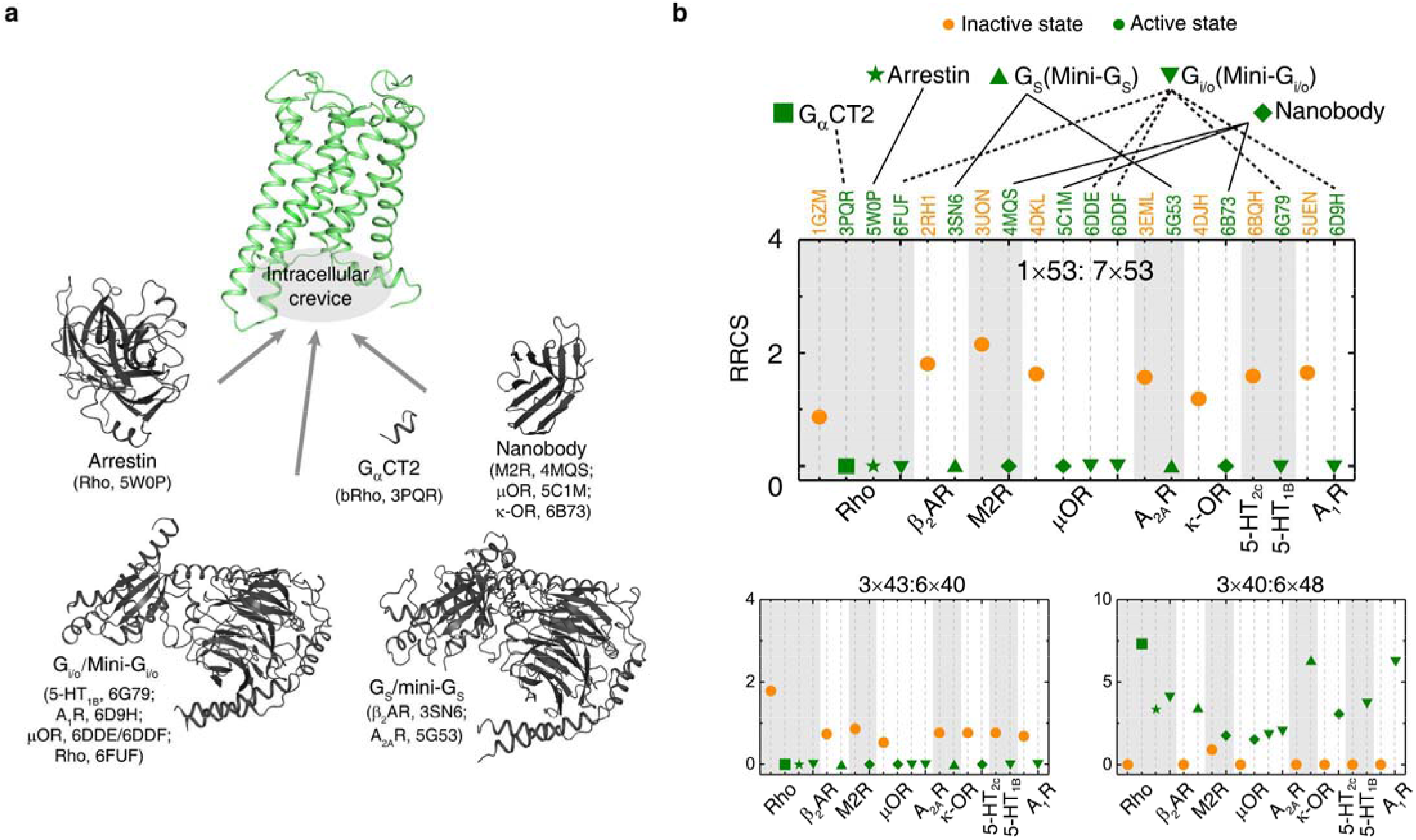
The universal activation pathway is conserved, regardless of the subtypes of intracellular effectors. **a**, Intracellular binding partners used in the active state structures. **b**, Comparison of RRCS for active (green) and inactive (orange) states of 8 receptors with different intracellular binding partners, including three recently solved cryo-EM structures of G_i/o_-bound receptors^38^ (5-HT_1B_, A_1_R, μOR) whose resolution were low (usually ≥3.8 Å for the GPCR part). Nevertheless, almost all conserved residue rearrangements in the pathway can be observed from these cryo-EM structures. Three of 34 residues pairs were shown here, see Figure 4—figure supplements 1 and 2 for the remaining 31 residue pairs. The following figure supplements are available for figure 4: **Figure supplement 1.** The switching conformation change is conserved upon receptor activation, regardless of the subtypes of intracellular effectors. **Figure supplement 2.** The repacking conformation change is conserved upon receptor activation, regardless of the subtypes of intracellular effectors.

### Molecular insights into the key steps of the universal receptor activation pathway

Receptor activation is triggered by ligand binding and is characterised by movements of different transmembrane helices. How does ligand-induced receptor activation connect the different and highly conserved motifs, rewire residue contacts and result in the observed changes in transmembrane helices? To this end, we analysed the universal activation pathway in detail and mapped, where possible, how they influence helix packing, rotation and movement (Figure 3). A qualitative analysis suggests the presence of four layers of residues in the activation pathway linking the ligand binding residues to the G protein binding region.

#### Layer 1

We did not see a single ligand-residue contact that exhibits conserved rearrangement, which accurately reflects the diverse repertoire of ligands that bind GPCRs^2, 12, 34^ (Figure 3—figure supplement 1). Instead, as a first common step, binding of diverse extracellular agonist converges to trigger an identical alteration of the transmission switch (3×40, 5×51, 6×44 and 6×48)^1, 21^ and Na^+^ pocket (2×50, 3×39, 7×45 and 7×49)^19, 24, 33^. Specifically, the repacking of an intrahelical contact between residues at 6×48 and 6×44, together with the switching contacts of residue at 3×40 towards 6×48 and residue at 5×51 towards 6×44, contract the TM3-5-6 interface in this layer. This reorganization initializes the rotation of the cytoplasmic end of TM6. The collapse of Na^+^ pocket^19, 24, 32–33^ leads to a denser repacking of the four residues (2×50, 3×39, 7×45 and 7×49), initiating the movement of TM7 towards TM3.

#### Layer 2

In parallel with these movements, two residues (6×40 and 6×41) switch their contacts with residue at 3×43, and form new contacts instead with residues at 5×58 and 5×55, respectively. Residues at 3×43, 6×40 and 6×41 are mainly composed of hydrophobic amino acids and referred as hydrophobic lock^22, 28, 39^. Its opening loosens the packing of TM3-TM6 and facilitates the outward movement of the cytoplasmic end of TM6, which is necessary for receptor activation. Additionally, N^7×49^ develops contacts with residue at 3×43 from nothing, facilitating the movement of TM7 towards TM3.

#### Layer 3

Upon receptor activation, Y^7×53^ loses its interhelical contacts^8^ with residues at 1×53 and 8×50, and forms new contacts with residues at 3×43, 3×46 and R^3×50^, which were closely packed with residues in TM6. Thus, the switching of contacts by Y^7×53^ strengthens the packing of TM3-TM7, while the packing of TM3-TM6 is further loosened with the outward movement of TM6.

#### Layer 4

Finally, the restrains on R^3×50^, including more conserved, local intrahelical contacts with D(E)^3×49^ and less conserved ionic lock with D(E)^6×30^, are eliminated^11, 14, 37^ and R^3×50^ is released. Notably, the switching contacts between R^3×50^ and residue at 6×37 are essential for the release of R^3×50^, which breaks the remaining contacts between TM3 and TM6 in the cytoplasmic end and drives the outward movement of TM6. The rewired contacts of R^3×50^ and other G protein contacting positions (3×53, 3×54, 5×61 and 6×33) make the receptor competent to bind to G protein on the cytosolic side. Together, these findings demonstrate that the intrahelical/interhelical and switching/repacking contacts between residues is not only critical to reveal the continuous and modular nature of the activation pathway, but also to link residue-level changes to transmembrane helix-level changes in the receptor.

### Universal activation pathway induced changes in transmembrane helix packing in GPCRs

To capture the patterns in the global movements of transmembrane helices, 8 residue pairs were chosen to describe the interhelical contacts between the cytoplasmic end of TM3 and TM6 as well as TM3 and TM7 (Figure 5a). Analysis of the RRCS_TM3-TM7_ (X-axis) and RRCS_TM3-TM6_ (Y-axis) for each of the 234 class A GPCR structures revealed distinct compact clusters of inactive and active states. Surprisingly, the inactive state has zero or close to zero RRCS_TM3-TM7_ regardless of the wide distribution of RRCS_TM3-TM6_. In contrast, the active state has a high RRCS_TM3-TM7_ and strictly zero RRCS_TM3-TM6_. Thus, receptor activation from inactive to active state occurs as a harmonious process of interhelical contacts changes: elimination of TM3-TM6 contacts, formation of TM3-TM7 contacts and repacking of TM5-TM6 (Figure 5b and Figure 5—figure supplement 1). In terms of global conformational changes, the binding of diverse agonists converges to trigger outward movement of the cytoplasmic end of TM6 and inward movement of TM7 towards TM3^8, 20, 23^, thereby creating an intracellular crevice for G protein coupling (Figure 5b).

**Figure 5.**
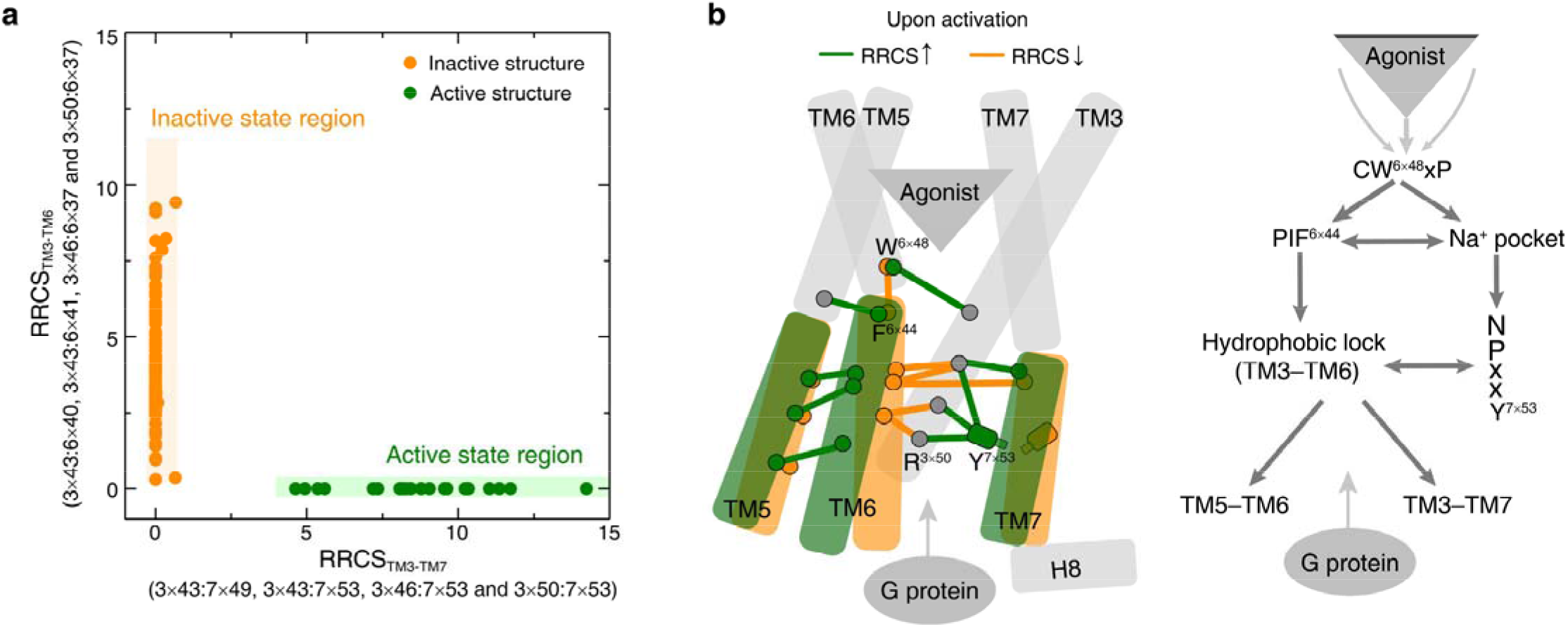
Universal activation model of class A GPCRs reveals the major changes upon GPCR activation. **a**, Active and inactive state structures form compact clusters in the 2D interhelical contact space: RRCS_TM3-TM7_ (X-axis) and RRCS_TM3-TM6_ (Y-axis). GPCR activation is best described by the outward movement of TM6 and inward movement of TM7, resulting in switch in the contacts of TM3 from TM6 to TM7. **b**, Universal activation model for class A GPCRs. Residues are shown in circles, conserved contact rearrangements of residue pairs upon activation are denoted by lines. The following figure supplement is available for figure 5: **Figure supplement 1.** Global conformational change upon activation.

### Experimental validation for the modular nature of the universal activation pathway

Based on the knowledge of the universal activation pathway, one would expect that mutations of residues in the pathway are likely to severely affect receptor activation. The two extreme consequences are constitutive activation (without agonist binding) or inactivation (abolished signalling). To experimentally test this hypothesis, we systematically designed site-directed mutagenesis for residues in the pathway on a prototypical receptor A_2A_R, aiming to create constitutively activating/inactivating mutations (CAM/CIM), by promoting/blocking residue and helix level conformational changes revealed in the pathway. 6/15 designed CAMs and 15/20 designed CIMs were validated by functional cAMP accumulation assays, and none of them were reported before for A_2A_R (Figure 6, Figure 6— figure supplement 1 and Figure 6—source data 1). The design of functional active/inactive mutants has been very challenging. However, the knowledge of universal activation pathway of GPCRs presented here greatly improves the success rate. The mechanistic interpretation of 21 successful predicted mutants is explained as below. We discuss the 14 unsuccessful predictions in the Figure 6—source data 2.

**Figure 6.**
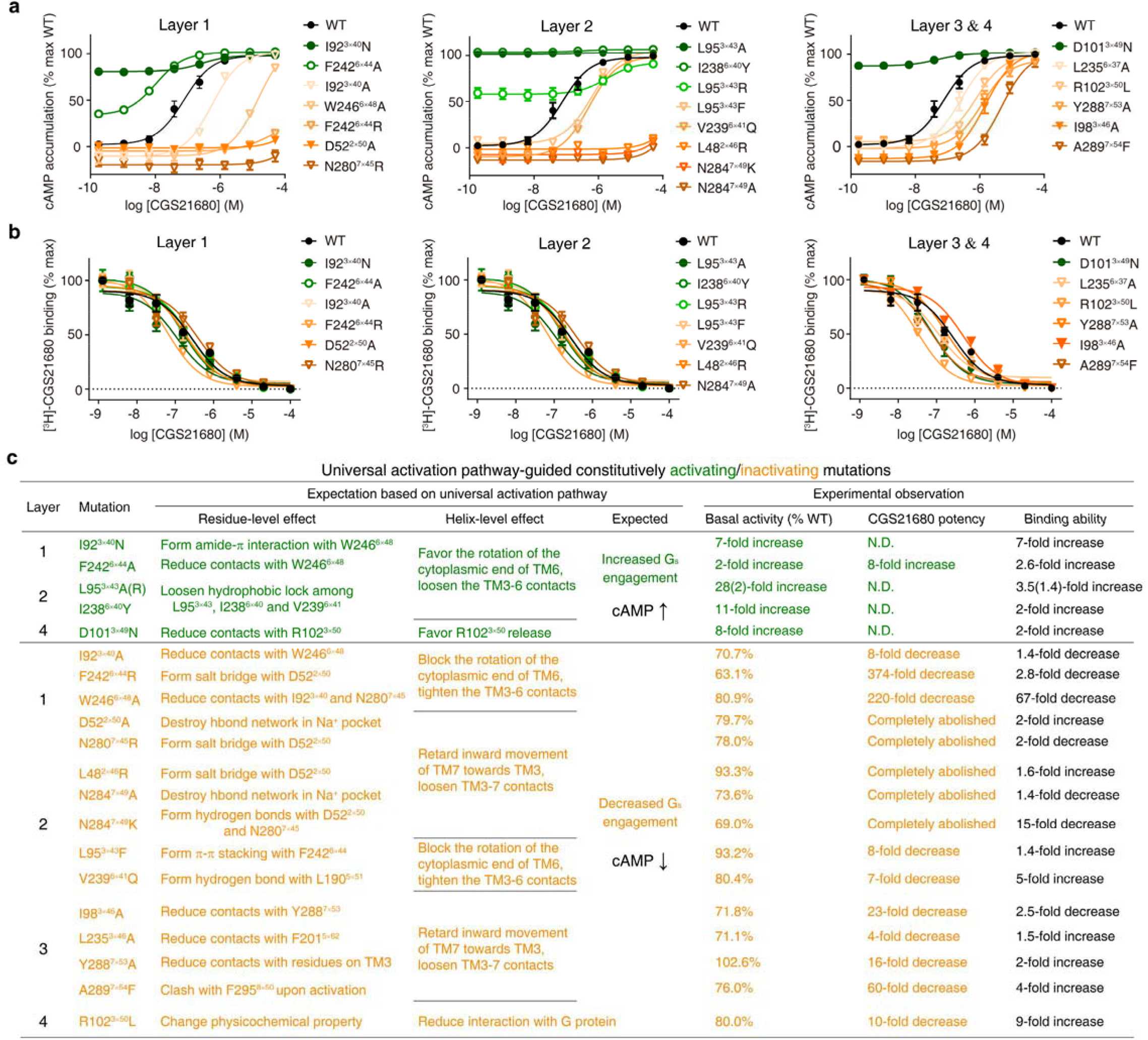
Experimental validation of the universal activation mechanism. **a**, cAMP accumulation assay and **b**, radioligand binding assay, validated the universal activation pathway-guided design of CAMs/CIMs for A_2A_R. WT, CAMs and CIMs are shown in black, green and orange, respectively. **c**, Mechanistic interpretation of universal activation pathway-guided CAMs/CIMs design. N.D.: basal activity was too high to determine an accurate EC_50_ value. The following data and figure supplement are available for figure 6: **Source data 1.** Functional and ligand binding properties of A2AR mutations. **Source data 2.** Analysis on the 14 unsuccessful predictions of A2AR CAMs/CIMs. **Figure supplement 1.** Experimental validation of universal activation pathway-guided CAM/CIM design.

In layer 1, the mutation I92^3×40^N likely stabilizes the active state by forming amide-π interactions with W246^6×48^ and hydrogen bond with the backbone of C185^5×461^, which rewires the packing at the transmission switch and initiates the outward movement of the cytoplasmic end of TM6; this mutation elevated the basal cAMP level by 7-fold. Conversely, I92^3×40^A would reduce the favourable contacts with W^6×48^ upon activation, which retards the initiation of the outward movement of TM6; this mutation resulted in a decrease in both basal cAMP level [71% of wild-type (WT)] and agonist potency (8-fold). Another example is the residue at 6×44, the mutation F242^6×44^R would stabilize the inactive state by forming salt bridge with D52^2×50^, which blocks the rotation of TM6 and thus abolishes G_s_ coupling; indeed this mutation greatly reduced basal cAMP level (to 63% WT) and agonist potency (by 374-fold). In contrast, F242^6×44^A would reduce contacts with W246^6×48^, loosen TM3-TM6 contacts, diminish the energy barrier of TM6 release and make outward movement of TM6 easier; consistently this mutation elevated the basal cAMP level (by 2-fold) and increased the agonist potency (by 8-fold). Mutations of residues forming the Na^+^ pocket, such as D52^2×50^A and N280^7×45^R, would destroy the hydrogen bond network at the Na^+^ pocket and retard the initiation of the inward movement of TM7. These mutations completely abolished agonist potency and greatly reduced the basal cAMP level (to 80% and 78% of WT, respectively).

In layer 2, the mutations L95^3×43^A/R and I238^6×40^Y would loosen the hydrophobic lock, weaken TM3-TM6 contacts, promote the outward movement of cytoplasmic end of TM6 and eventually make receptor constitutively active; this is reflected by remarkably high basal cAMP production (28-, 2- and 11-fold increase, respectively). Notably, mutations at/near the Na^+^ pocket, L48^2×46^R and N284^7×49^K, could lock the Na^+^ pocket at inactive packing mode by introducing salt bridge with D52^2×50^, thus block the inward movement of TM7 towards TM3. As expected, these mutations completely abolished agonist potency. The CIMs at/near the Na^+^ pocket (from both layer 1 and 2) reflect the subtle inward movement of TM7 towards TM3 is essential for receptor activation, which is often underappreciated and overshadowed by the movement of TM6. In line with this, two mutations on TM7, N284^7×49^A and Y288^7×53^A, attenuate the TM3-TM7 contacts upon activation and completely abolished or greatly reduced (by 16-fold) agonist potency, respectively.

In layer 3, I98^3×46^A likely reduces contacts with Y288^7×53^, weakens the packing between TM3-TM7, and retards the movement of TM7 towards TM3; similarly, L235^6×37^A would reduce contacts with F201^5×62^, weaken the packing between TM5-TM6, and makes the TM6 movement towards TM5 more difficult. In line with the interpretation, these mutations resulted in reduced basal cAMP level (72% and 71% WT, respectively) and decreased agonist potency (23- and 4-fold, respectively). These results are consistent with previous findings on vasopressin type-2 receptor (V2R)^8^. In layer 4, D101^3×49^N likely diminishes its intrahelical interactions with R102^3×50^ and thus makes the release of the latter easier, which in turn promotes the G protein recruitment. Consistent with this possibility, this mutation led to a greatly elevated basal cAMP level (8-fold).

Despite these A_2A_R mutants greatly affect receptor activation, our radioligand binding assay shows that they generally retain the agonist binding ability, with the exception of two CIMs: W246^6×48^A and N284^7×45^K (Figure 6b, c and Figure 6—source data 1). This suggests that the universal activation pathway is modular and that such an organization allows for a significant number of residues involved in agonist binding to be uncoupled from receptor activation/inactivation and G protein binding.

### The universal activation pathway allows mechanistic interpretation of mutations

Four hundred thirty five disease-associated mutations were collected, among which 28% can be mapped to the universal activation pathway, much higher than that to the ligand-binding and G protein-binding regions (20% and 7%, respectively) (Figure 7a, b). Furthermore, 272 CAMs/CIMs from 41 receptors (Figure 7c) were mined from the literature for the 14 hub residues (*i.e.*, residues that have more than one edges in the pathway).

**Figure 7.**
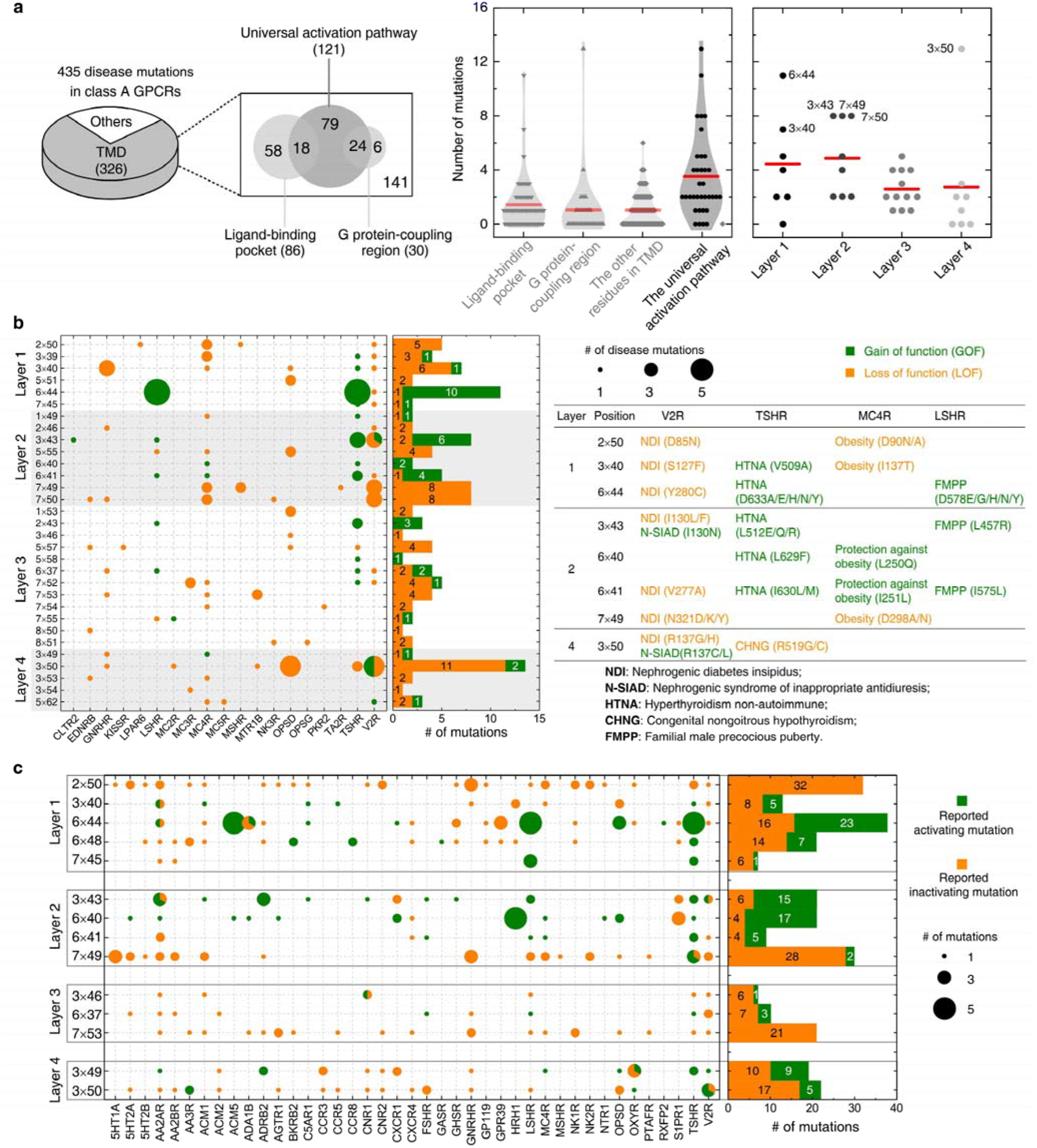
Importance of the universal activation pathway in pathophysiological and biological context. **a**, Comparison of disease-associated mutations in the universal activation pathway (further decomposed into layers 1-4), ligand-binding pocket, G protein-coupling region and other regions. Red line denotes the mean value. **b**, Mapping of disease-associated mutations in class A GPCRs to the universal activation pathway. **c**, Key roles of the residues constituting the universal activation pathway have been reported in numerous experimental studies on class A GPCRs. 272 CAMs/CIMs from 41 receptors were mined from the literature for the 14 hub residues (*i.e.*, residues that have more than one edges in the pathway). The following data and figure supplements are available for figure 7: **Source data 1.** Constitutively activating/inactivating mutations for the 14 hub residues in the universal activation pathway **Figure supplement 1.** The universal activation pathway can be used to mechanistically interpret disease-associated mutations and CAMs/CIMs. **Figure supplement 2.** Residues in the universal activation pathway are more conserved than other functional regions of GPCR.

The average number of disease-associated mutations in the universal activation pathway is much higher than that of ligand-binding pocket, G protein-binding pocket, and residues in other regions (2.5-, 3.5- and 3.5-fold, respectively), reflecting the enrichment of disease-associated mutations on the pathway (Figure 7a). Within the universal activation pathway, the enrichment of disease mutations and CAMs/CIMs in layers 1 and 2 is noteworthy, which highlights the importance of signal initiation and hydrophobic lock opening, and further supports the modular and hierarchical nature of GPCR activation (Figures 3 and 5b). Notably, for certain residues, such as D^2×50^ and Y^7×53^, only loss-of-function disease mutations or CIMs were observed (Figures 7 and 8b), implying they are indispensable for receptor activation and the essential role of TM7 movement (Figures 3 and 5).

The functional consequence of these single point mutations can be rationalized by analysing if they are stabilizing/destabilizing the contacts in the universal activation pathway or promoting/retarding the required helix movement upon activation (Figure 7b and Figure 7—figure supplement 1). For example, I130^3×43^N/F (in layer 2 of the universal activation pathway) in V2R was reported as a gain-/loss-of-function mutation that causes nephrogenic syndrome of inappropriate antidiuresis^40^ or nephrogenic diabetes insipidus^41^, respectively. I130^3×43^N/F likely loosens/stabilizes the hydrophobic lock, weakens/strengthens the TM3-TM6 packing and leads to constitutively active/inactive receptors. Another example is T58^1×53^M in rhodopsin, which was reported as a loss-of-function mutation that causes retinitis pigmentosa 4^42^. T58^1×53^M likely increases hydrophobic contacts with Y306^7×53^ and P303^7×50^, which retards the inward movement of TM7 towards TM3 and eventually decreases G protein recruitment. As in the case of disease-associated mutations, CAMs/CIMs that have been previously reported in the literature can also be interpreted by the framework of universal activation pathway (Figure 7—figure supplement 1b). For example, F248^6×44^Y in CXCR4^28^ was reported as a CIM. This residue likely forms hydrogen bond with S123^3×39^, which blocks the rotation of the cytoplasmic end of TM6, and decreases G protein engagement.

Not surprisingly, the 35 residues constituting the pathway are highly conserved across class A GPCRs, dominated by physiochemically similar amino acids (Figure 7—figure supplement 2). The average sequence similarity of these positions across 286 non-olfactory class A receptors is 66.2%, significantly higher than that of ligand-binding pockets (31.9%) or signalling protein-coupling regions (35.1%). Together, these observations suggest that the modular and hierarchical nature of the activation pathway allows decoupling of the ligand-binding pocket, G protein-binding pocket and the residues contributing to the universal activation mechanism. Such an organization of the receptor might facilitate the uneven sequence conservation between different regions of GPCRs, confers their functional diversity in ligand recognition and G protein binding while still retaining a common activation mechanism.

## Discussion

Using a novel, quantitative residue contact descriptor, RRCS, and a family-wide comparison across 234 structures from 45 class A GPCRs, we reveal a universal, modular activation pathway that directly links ligand-binding pocket and G protein-coupling regions. Key residues that connect the different modules allows for the decoupling of a large number of residues in the ligand-binding site, G protein contacting region and residues involved in the activation pathway. Such an organization may have facilitated the rapid expansion of GPCRs through duplication and divergence, allowing them to evolve independently and bind to diverse ligands due to removal of the constraint (i.e. between a large number of ligand binding residues and those involved in receptor activation). This model unifies many previous important motifs and observations on GPCR activation in the literature (CWxP^11, 20, 33^, DRY^11, 14, 37^, Na^+^ pocket^19, 24, 33^, NPxxY^20, 23^, PIF^3, 36^ and hydrophobic lock^22, 28, 39^] and is consistent with numerous experimental findings^21–22, 28, 33, 39^.

We focused on the universal activation pathway (*i.e.*, the common part of activation mechanism shared by all class A GPCRs and various intracellular effectors) in this study. Obviously, individual class A receptor naturally has its intrinsic activation mechanism(s), as a result of the diversified sequences, ligands and physiological functions. Indeed, receptor-specific activation pathways (including mechanisms of orthosteric, positive or negative allosteric modulators, biased signalling/selectivity of downstream effectors)^5, 9, 17, 43–48^ have been revealed by both experimental studies including biophysical (such as X-ray, cryo-EM, NMR, FRET/BRET, DEER)^2, 9, 14, 27, 33, 43, 49–52^, biochemical^28, 39^ and computational approaches (such as evolutionary trace analysis^26, 30^ and molecular dynamics simulations^16, 25, 31, 53^), especially for the prototypical receptors such as rhodopsin, β_2_-adrenergic and A_2A_ receptors. These studies demonstrated the complexity and plasticity of signal transduction of GPCRs. The computational framework we have developed may assist us in better understanding the mechanism of allosteric modulators, G protein selectivity and diverse activation processes via intermediate states as more GPCR structures become available. While we interpret the changes as a linear set of events, future studies aimed at understanding dynamics could provide further insights into how the common activation mechanism operates in individual receptors.

Given the universal and conserved nature of the pathway, we envision that the knowledge of the common activation pathway can not only be used to mechanistically interpret the effect of mutations in biological and pathophysiological context^54^ but also to rationally introduce mutations in other receptors by promoting/blocking residue and helix level movements that are essential for activation. Such protein engineering approaches can obtain receptors in specific conformational states to accelerate structure determination studies using X-ray crystallography or electron microscopy in future. The approach developed here can be readily adapted to map allosteric pathways and reveal mechanisms of action for other key biological systems such as kinases, ion channels and transcription factors.

---

**Figure 1—figure supplement 1.**
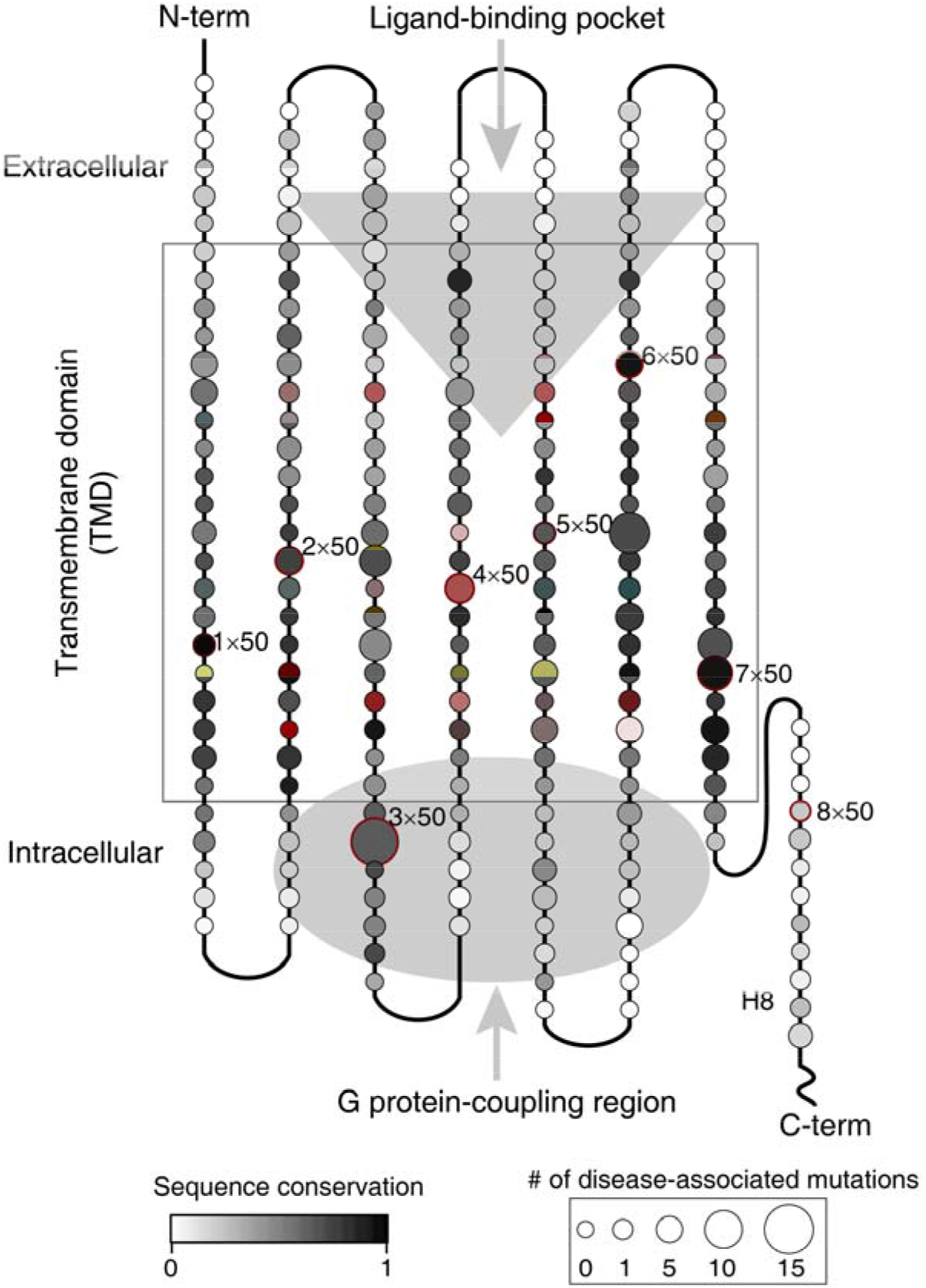
The pattern of conservation of residues and the map of number of disease-associated mutations on human class A GPCRs. The alignment of 286 non-olfactory, class A human GPCRs were obtained from the GPCRdb^35, 54–55^ and sent for the sequence conservation score calculation for all residue positions by the Protein Residue Conservation Prediction^56^ tool with scoring method “property entropy”^57^. To obtain disease-associated mutations, we performed database integration and literature investigation for all 286 non-olfactory class A GPCRs. Four commonly used databases (UniProt^58^, OMIM^59^, Ensembl^60^ and GPCRdb^54–55^) were first filtered by disease mutations and then merged. Finally, we collected 435 disease mutations from 61 class A GPCRs (Figure 1—Source data 2).

**Figure 2—figure supplement 1.**
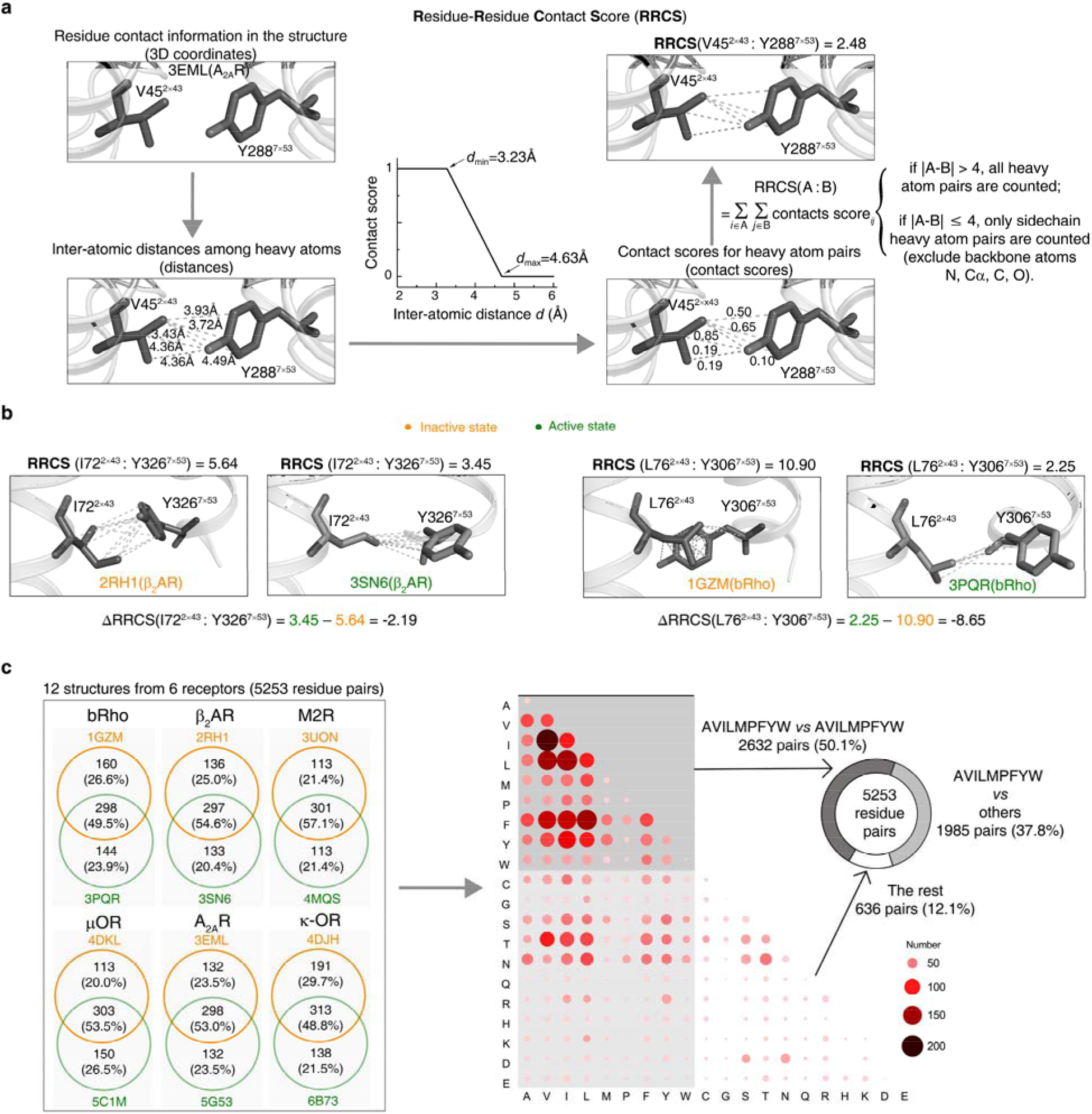
Calculation of RRCS and ΔRRCS. **a**, Workflow of RRCS calculation. **b**, Examples of RRCS and ΔRRCS calculation for two residues pairs. **c**, Statistics of residue contacts and contact types for six receptors (bRho, β_2_AR, M2R, μOR, A_2A_R and κ-OR) in their inactive and active states. Contact type describes physicochemical properties of two interacted amino acids that form a pair. The amino acids with hydrophobic side chains (one-letter code: A, V, I, L, M, P, F, Y, W) contribute to the majority of residue contacts, either within themselves (50.1%) or with other amino acids (37.8%).

**Figure 3—figure supplement 1.**
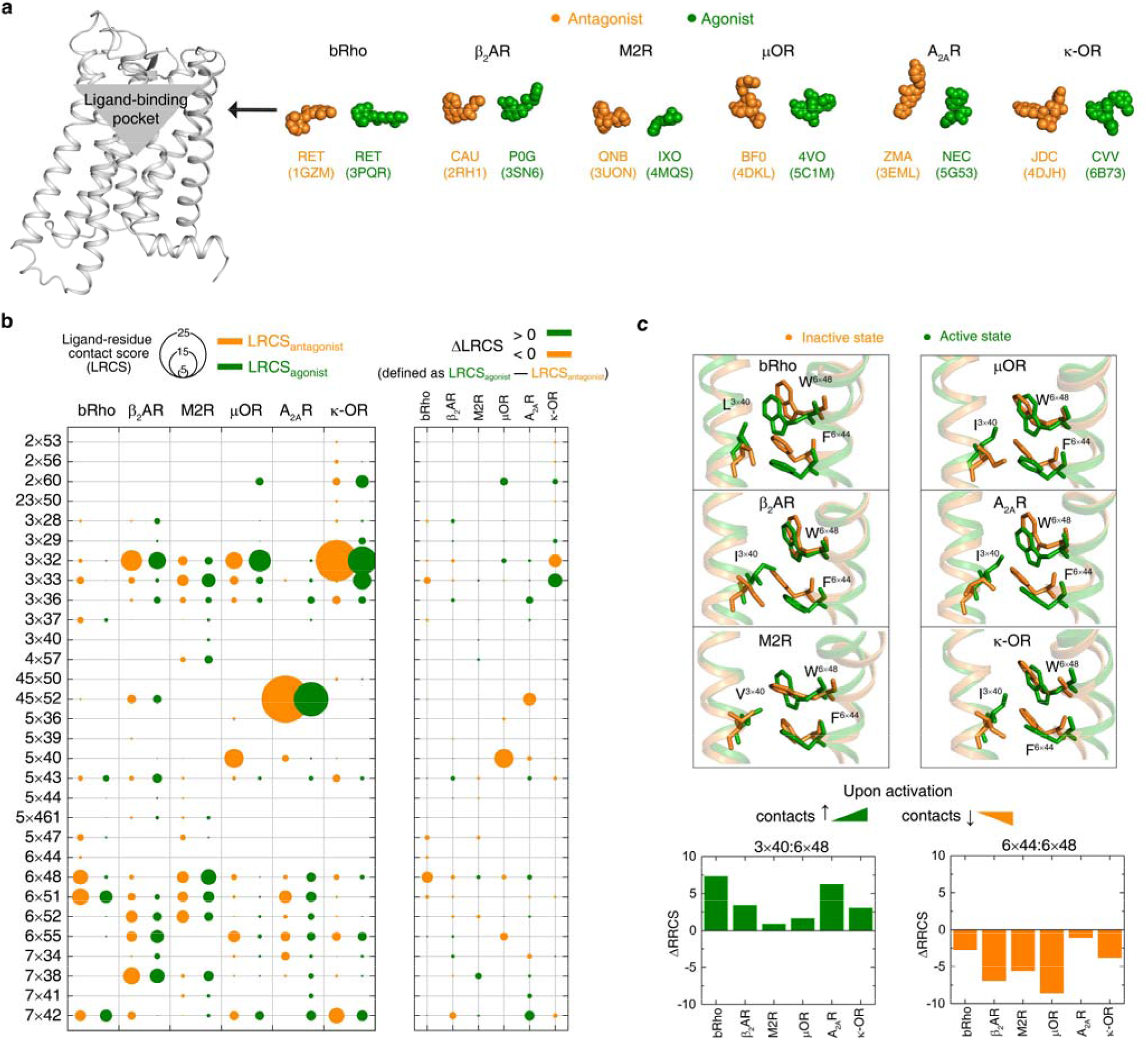
Rearrangements of ligand-residue contacts in ligand-binding pocket are not conserved, reflecting diverse ligand recognition modes. **a**, Sphere representation of antagonist- and agonist-bound receptor crystal structures. **b**, Diverse LRCS and ΔLRCS reveal the repertoire of ligand recognition across class A GPCRs. The agonist or antagonist was treated as a single residue when calculating LRCS and ΔLRCS. As shown by the calculated ΔRRCS, no ligand-residue pair exhibits conserved rearrangements upon activation. **c**, Conserved conformational changes were only observed at the very bottom of ligand-binding pocket (6×48, 3×40 and 6×44).

**Figure 4—figure supplement 1.**
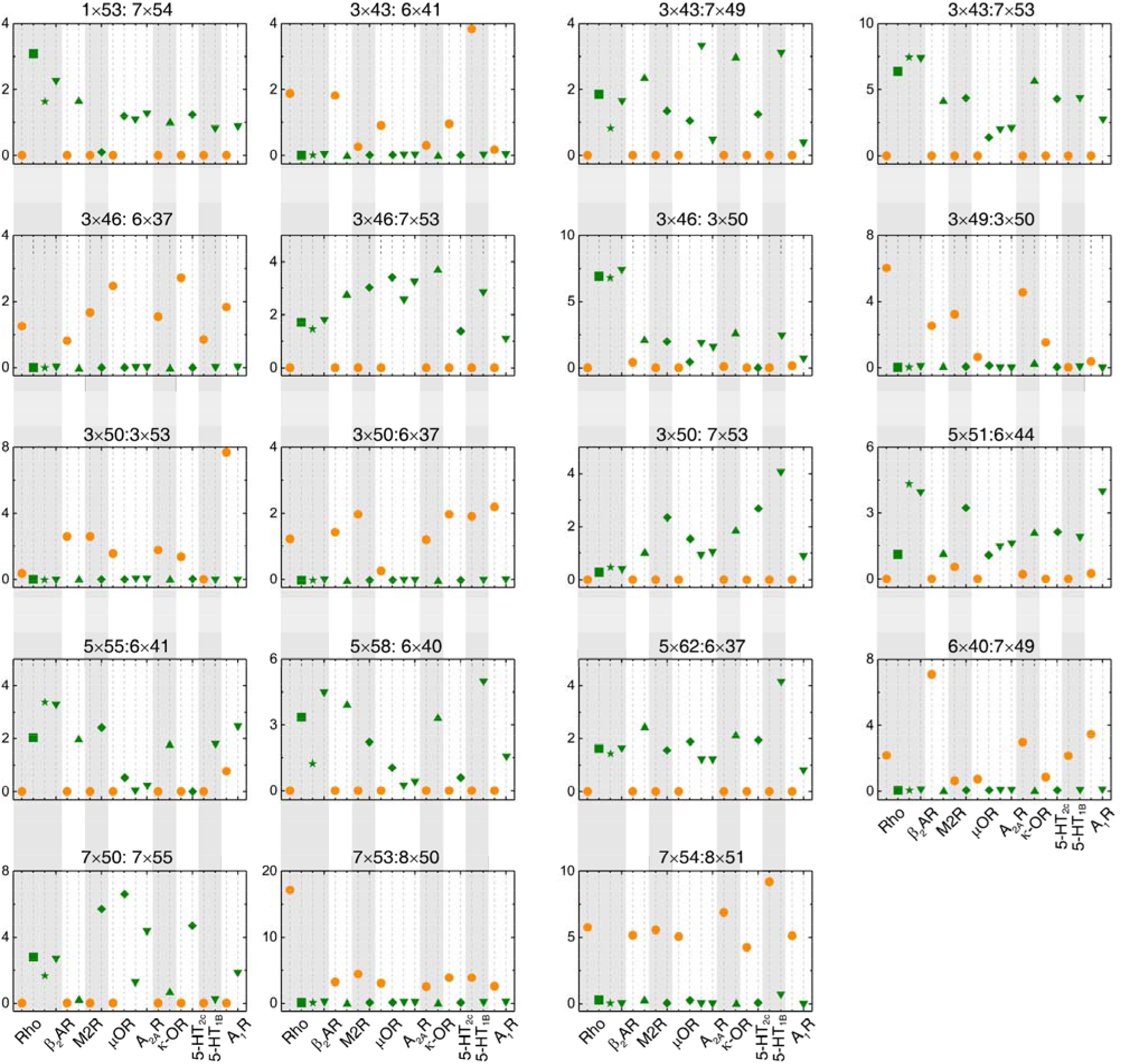
The switching conformation change is conserved upon receptor activation, regardless of the subtypes of intracellular effectors. Comparison of RRCS for active (green) and inactive (orange) states of 8 receptors with different intracellular binding partners, including three recently solved cryo-EM structures of G_i/o_-bound receptors (5-HT_1B_, A_1_R, μOR) whose resolution were low (usually ≥3.8 Å for the GPCR part)^38^. Nevertheless, almost all conserved residue rearrangements in the pathway can be observed from these cryo-EM structures. Nineteen of 34 residues pairs were shown here, see Figure 4 and Figure 4-figure supplement 2 for the remaining residue pairs.

**Figure 4—figure supplement 2.**
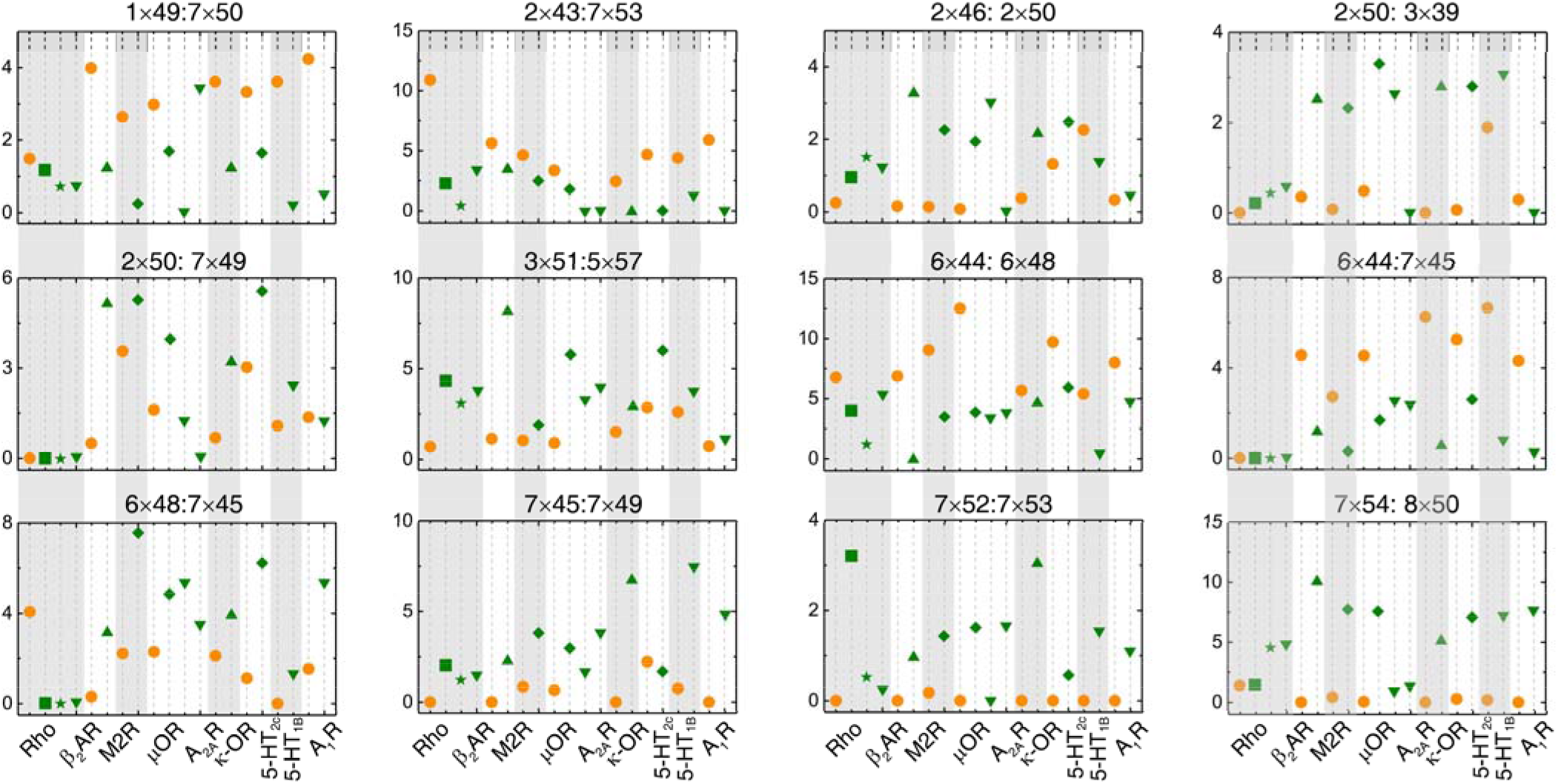
The repacking conformation change is conserved upon receptor activation, regardless of the subtypes of intracellular effectors. Comparison of RRCS for active (green) and inactive (orange) states of 8 receptors with different intracellular binding partners, including three recently solved cryo-EM structures of G_i/o_-bound receptors (5-HT_1B_, A_1_R, μOR) whose resolution were low (usually ≥3.8 Å for the GPCR part)^38^. Nevertheless, almost all conserved residue rearrangements in the pathway can be observed from these cryo-EM structures. Twelve of 34 residues pairs were shown here, see Figure 4 and Figure 4-figure supplement 1 for the remaining residue pairs.

**Figure 5—figure supplement 1.**
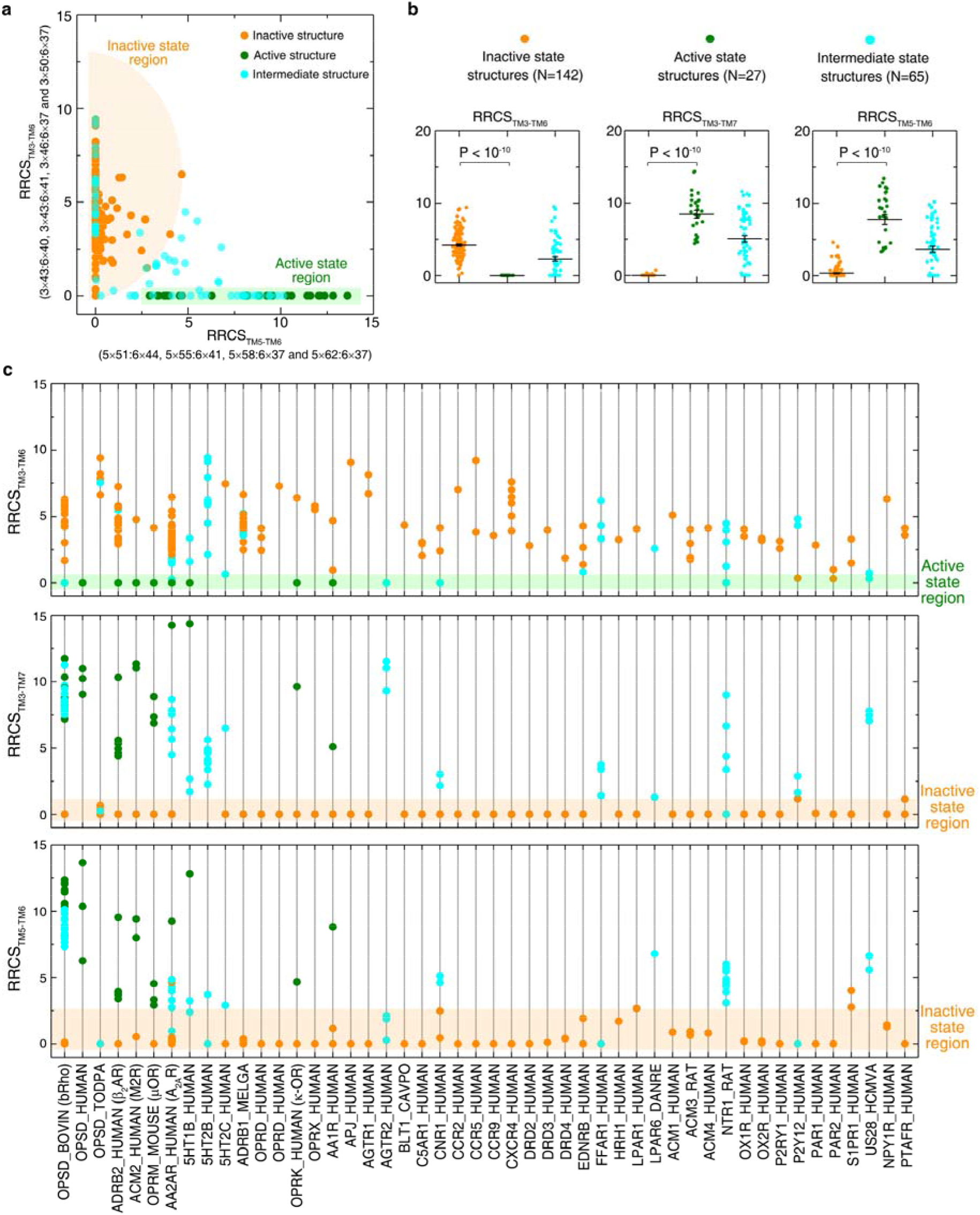
Global conformational change upon activation. **a**, Distinct clustering of inactive- and active-state structures in 2-dimentional interhelical contact space RRCS_TM5-TM6_ *vs*. RRCS_TM3-TM6_. **b**, The interhelical contacts comparison between inactive- and active-state structures. **c**, Receptor-specific interhelical contacts for all class A GPCR structures (inactive, intermediate and active states are coloured in orange, cyan and green, respectively). These results demonstrate that receptor activation involves the elimination of TM3-TM6 contacts, formation of TM3-TM7 and TM5-TM6 contacts, reflecting the outward movement of the cytoplasmic end of TM6 away from TM3, the inward movement of TM7 towards TM3 and the repacking of TM5 and TM6.

**Figure 6—source data 1.**
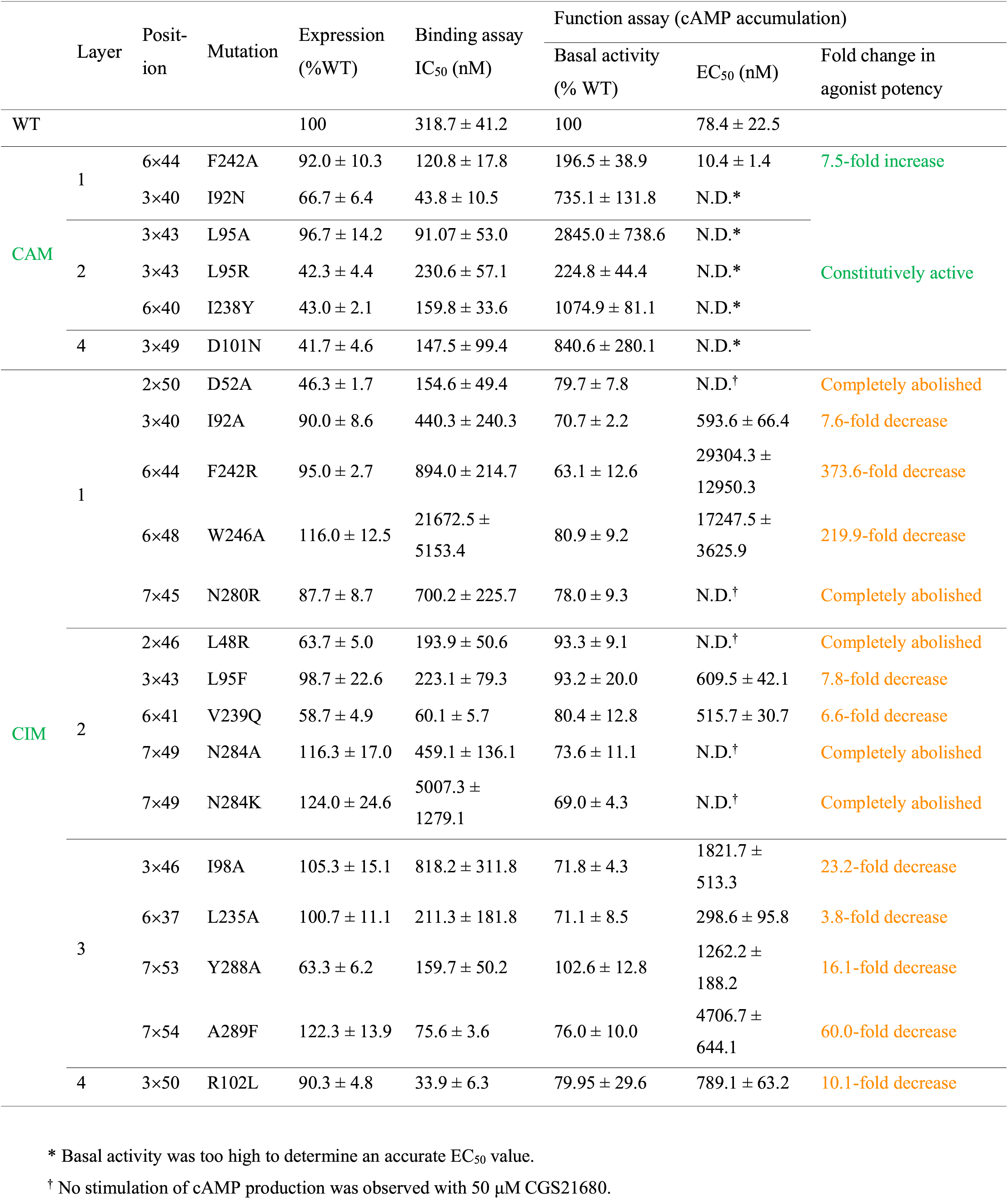
Functional and ligand binding properties 409 of A2AR mutations.

**Figure 6—source data 2.**
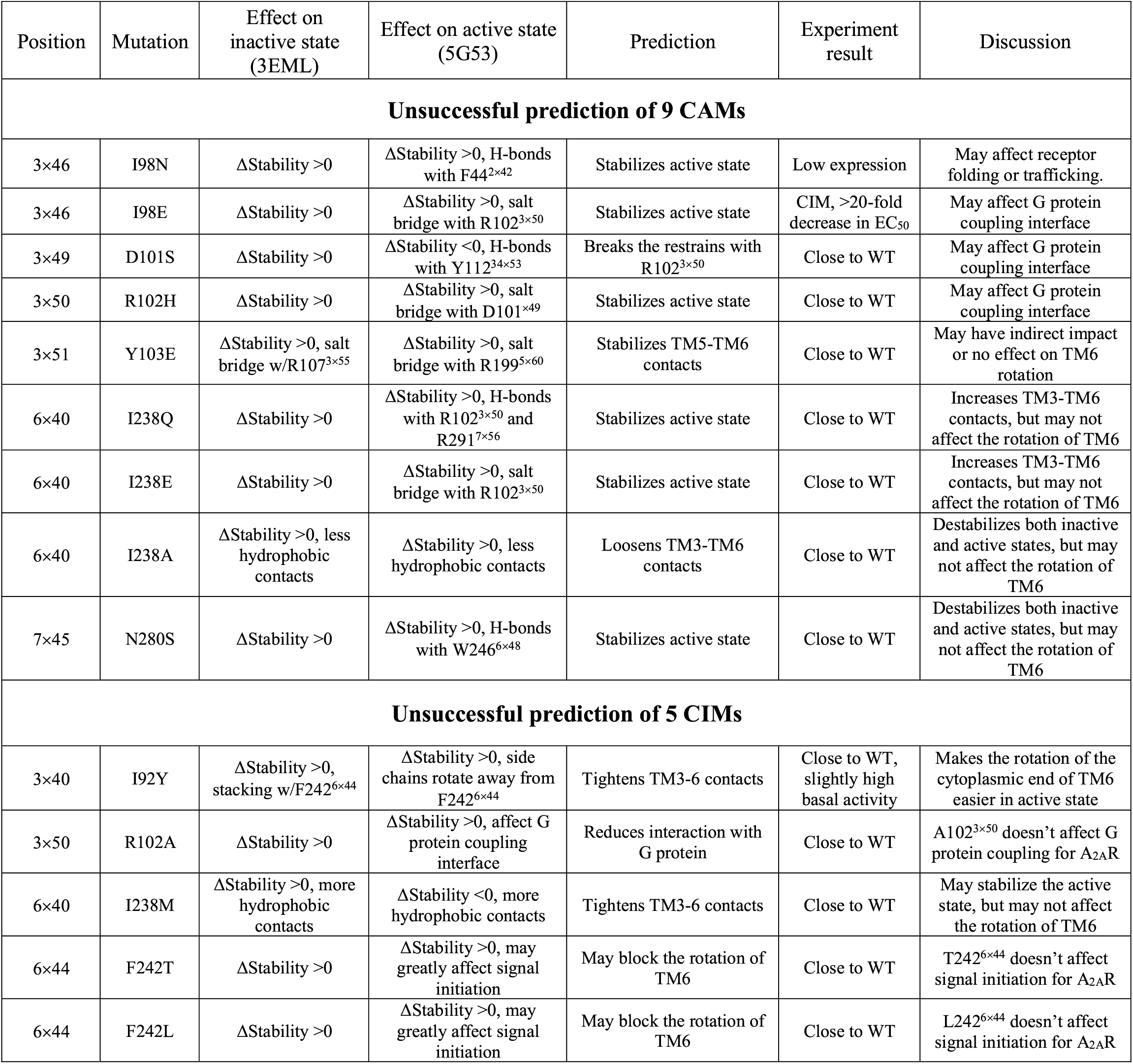
Analysis on the 14 unsuccessful predictions 413 of A2AR CAMs/CIMs. ΔStability (>0 means destabilized; <0 means stabilized) is the change of receptor stability when a mutation was introduced, calculated by Residue Scanning module in BioLuminate^61^. WT, wild-type.

**Figure 6—figure supplement 1.**
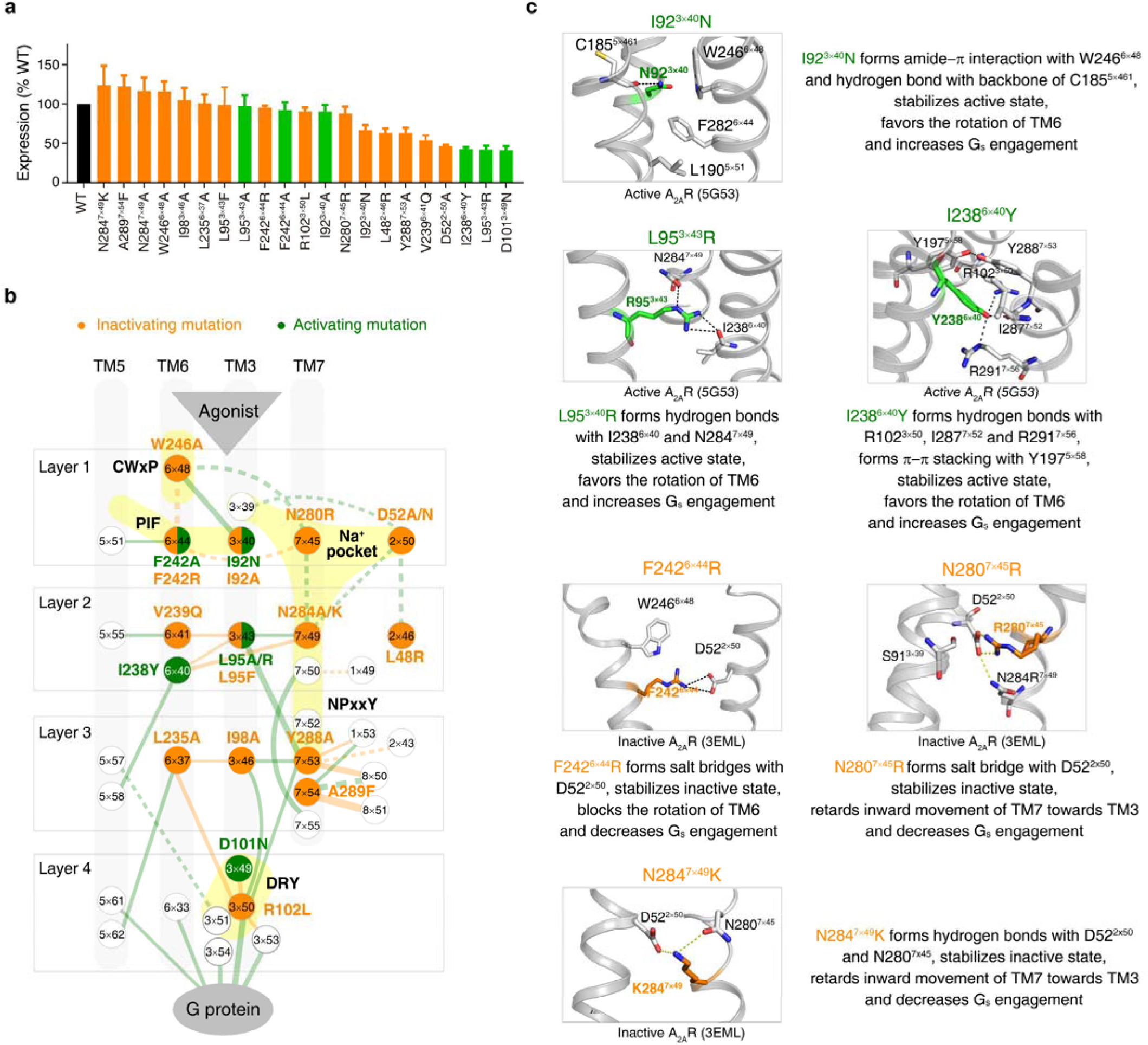
Experimental validation of universal activation pathway-guided CAM/CIM design. **a**, Cell surface expression of the WT A_2A_R and its mutants. WT, CAMs and CIMs are coloured by black, orange and green, respectively. **b**, Mapping of validated CAMs/CIMs to the universal activation pathway. **c**, The mechanisms of CAM/CIM design. CAMs and CIMs are in green and orange, respectively.

**Figure 7—figure supplement 1.**
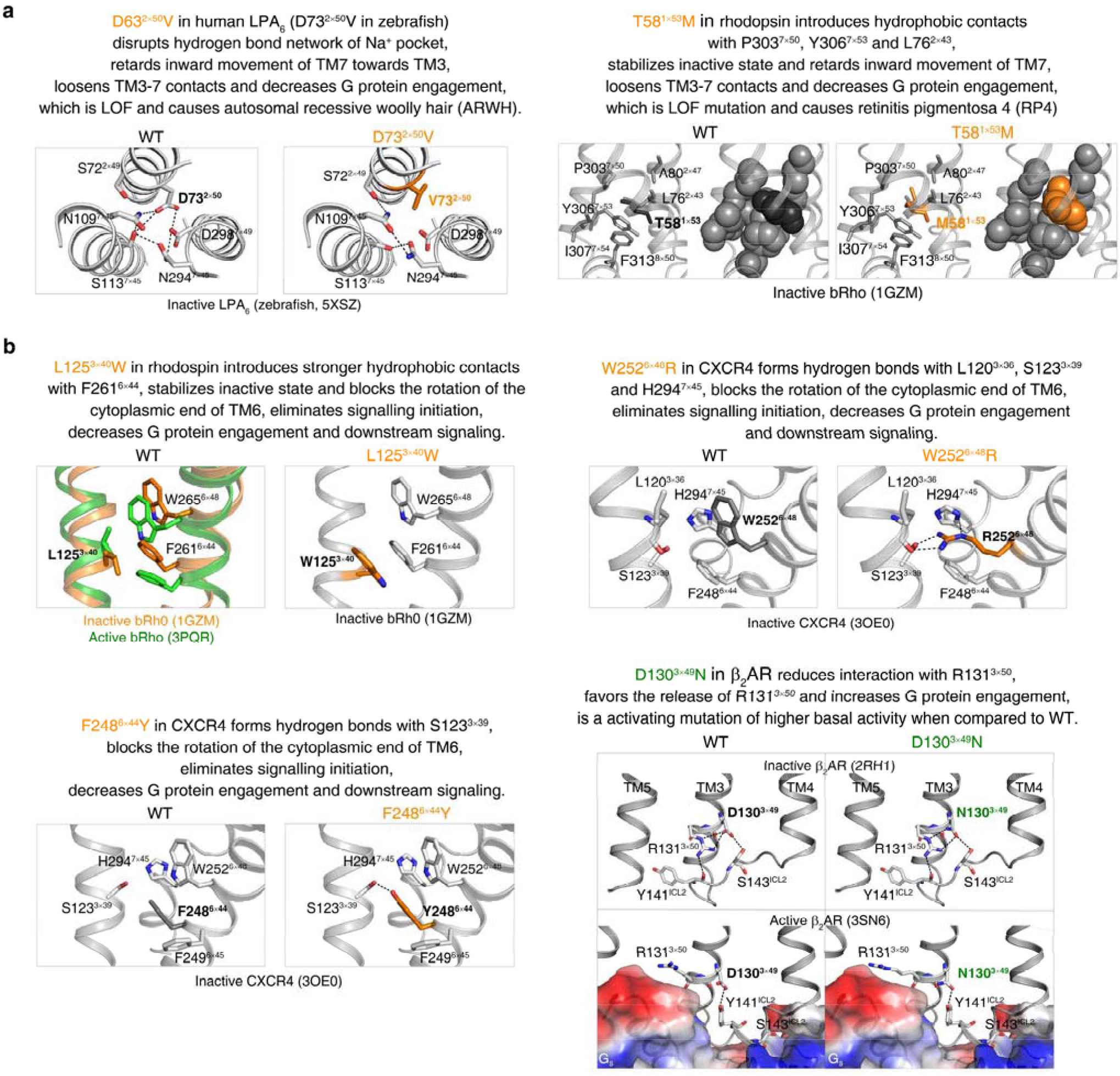
The universal activation pathway can be used to mechanistically interpret disease-associated mutations and CAMs/CIMs. **a**, Pathway-guided mechanistic interpretations of two disease mutations. **b**, Pathway-guided mechanistic interpretations of four CAMs/CIMs.

**Figure 7—figure supplement 2.**
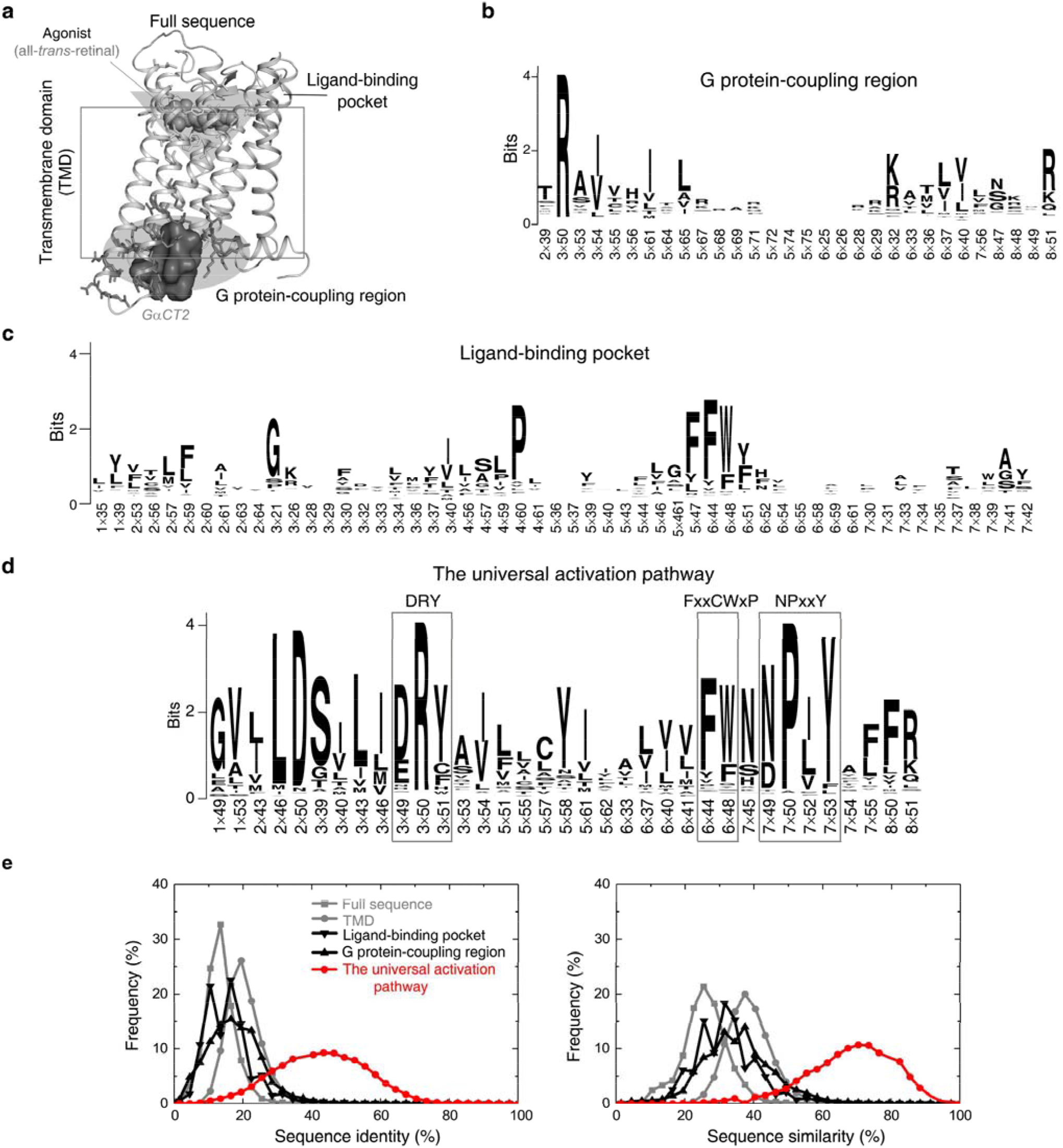
Residues in the universal activation pathway are more conserved than other functional regions of GPCR. **a**, Illustration of different functional regions of GPCR. **b-d**, Sequence pattern of the G protein-coupling region (**b**), ligand-binding pocket (**c**) and the universal activation pathway (**d**). **e**, Distribution of sequence identity (left) and similarity (right) for functional regions across 286 non-olfactory class A receptors.

## Materials and Methods

### Glossary

Transmembrane domains (TMD): the core domain exists in all GPCRs, and consists of seven-transmembrane helices (TM1-7) that are linked by three extracellular loops (ECL1-3) and three intracellular loops (ICL1-3).

GPCRdb numbering scheme: a structure-based numbering system for GPCRs^35, 62^, an improved version of sequence-based Ballesteros-Weinstein numbering^63^ that considers structural distortions such as helical bulges or constrictions. The most conserved residue in a helix n is designated n×50, while other residues on the helix are numbered relative to this position.

Node: a point in a network at which lines intersect, branch or terminate. In this case, nodes represent amino acid residues.

Edge: a connection between the nodes in a network. In this case, an edge represents a residue-residue contact.

Hub: a node with two or more edges in a network.

Constitutively activating mutation (CAM): a mutant that could increase the inherent basal activity of the receptor by activating the G protein-signalling cascade in the absence of agonist.

Constitutively inactivating mutation (CIM): a mutant completely abolishes receptor signalling.

### GPCR structure data set

As of October 1, 2018, there are 234 released structures of 45 class A GPCRs with resolution better than 3.8 Å (Figure 1—Source data 1), which covers 71% (203 out of 286 receptors, including 158 receptors that have no structures but share >50% sequence similarity in the TMD with the 45 structure-determined receptors) of class A GPCRs (Figure 1a). Based on the type of bound ligand and effector, these structures could be classified into three states: inactive state (antagonist or inverse agonist-bound, 142 structures from 38 receptors), active state (both agonist- and G protein/G protein mimetic-bound, 27 structures from 8 receptors) and intermediate state (only agonist-bound, 65 structures from 15 receptors). In this study, we primarily focused on conformational comparison between inactive- and active- state structures, while also investigating the intermediate state structures. In the structure data set, 7 receptors have both inactive and active structures: rhodopsin (bRho), β_2_-adrenergic receptor (β_2_AR), M2 muscarinic receptor (M2R), μ-opioid receptor (μOR), adenosine A_2A_ receptor (A_2A_R), κ-opioid receptor (κ-OR) and adenosine A_1A_ receptor (A_1A_R), the active state structure of which was recently determined by cryo-EM. In addition, 32 receptors have either inactive or active structures (Figure 1—Source data 1).

### Calculation of residue-residue contact score (RRCS)

We developed a much finer distance-based method (than coarse-grained Boolean descriptors such as contact map and residues contact^64–66^), namely residue-residue contact score (RRCS). For a pair of residues, RRCS is calculated by summing up a plateau-linear-plateau form atomic contact score adopted from GPCR–CoINPocket^34, 67–69^ for each possible inter-residue heavy atom pairs (Figure 2—figure supplement 1a). GPCR–CoINPocket is a modified version of the hydrophobic term of ChemScore^64–65^ that has been successfully used to describe hydrophobic contribution to binding free energy between ligand and protein. RRCS can describe the strength of residue-residue contact quantitatively in a much more accurate manner than Boolean descriptors^8, 10^. For example, Boolean descriptors do not capture side chain repacking if the backbone atoms of the two residues are close to each other (e.g., translocation of Y^7×53^ away from residue at 2×43 upon GPCR activation) and local contacts involving adjacent residues (residues within four/six amino acids in protein sequence) (e.g., disengagement between D/E^3×49^ and R^3×50^), while both cases can be well reflected by the change of RRCS (Figure 2c and Figure 2—figure supplement 1b).

All RRCS data can be found in Figure 2—Source data 1. The computational details are listed as below:

i. For the residue pairs between adjacent residues that are within four amino acids in protein sequence, only side chain heavy atom pairs were considered, atom pairs involving in backbone atoms (Cα, C, O, N) were excluded, since the latter seldom change during GPCR activation. For other residue pairs, all possible heavy atom pairs (including backbone atoms) were included when calculating RRCS.
ii. Atomic contact scores are solely based on interatomic distance, and they were treated equally without weighting factors such as atom type or contact orientation. In principle, weighting of atomic contact by atom type and/or orientation would improve residue-residue contact score. However, parameterization of atom type or contact orientation is relatively arbitrary, subjective and complicated, especially considering the lipid bilayer environment surrounding GPCRs. Our preliminary study for twelve structures from six receptors (bRho, β_2_AR, M2R, μOR, A_2A_R and κ-OR) revealed that amino acids with hydrophobic side chains (one-letter code: A, V, I, L, M, P, F, Y, W) contribute to the majority (∼88%) of residue pairs. Meanwhile, ionic lock opening of well-known motif DRY upon receptor activation can be adequately reflected by RRCS change between D/E^3×49^ and R^3×50^. These results suggest that interatomic distance-dependent residue pair contact score may represent an acceptable approximation of actual (either hydrophobic or charge-charge) interaction energies^34^ and is accurate enough for identifying conserved rearrangements of residue contacts upon receptor activation.
iii. The quality of structures is extremely important for RRCS calculation. We adopted two criteria to exclude unreliable structures and residues: (a) crystal structures whose resolution is ≥3.8 Å. Structures in this category are: 5DGY (7.70 Å), 2I37 (4.20 Å), 2I36 (4.10 Å), 5TE5 (4.00 Å), 4GBR (4.00 Å), 5NJ6 (4.00 Å), 5V54 (3.90 Å), 2I35 (3.80 Å), 5D5B (3.80 Å), 4XT3 (3.80 Å); (b) residues whose residue-based real-space R-value (RSR^70^) is greater than 0.35. RSR is measure of how well ‘observed’ and calculated electron densities agree for a residue. RSR ranges from 0 (perfect match) to 1 (no match); RSR greater than 0.4 indicates a poor fit^71^. Here we adopted a stricter cut-off, 0.35. Among the 234 class A GPCR structures, 156 have available RSR information^72^ (http://eds.bmc.uu.se), with 8.8% residues have RSR >0.35 and they are omitted in our analysis. For the 35 residues that constitute the universal activation pathway, 255 out of 5460 RSR data points (∼4.7%, lower than 8.8% for all residues) were omitted for having RSR values >0.35.
iv. For structures with multiple chains, RRCS were the average over all chains. For residues with multiple alternative conformations, RRCS is the sum of individual values multiplied by the weighting factor: occupancy value extracted from PDB files. Small molecule/peptide ligand, or intracellular binding partner (G protein or its mimetic) was treated as a single residue.
v. For the family-wide comparison of conformational changes upon activation, structurally equivalent residues are numbered by GPCRdb numbering scheme^35, 62^. Of the 35 residues in the universal activation pathway, their GPCRdb numbering in all structures is almost identical to the Ballesteros– Weinstein numbering^63^, the exceptions are residues at 6×37, 6×41 and 6×44 for five receptors: FFAR1, P2Y_1_, P2Y_12_, F2R and PAR2, which are all from the delta branch of class A family.

### Identification of conserved rearrangements of residue contacts upon activation

Using RRCS, structural information of TMD and helix 8 in each structure can be decomposed into 400∼500 residue pairs with positive RRCS. ΔRRCS, defined as RRCS_active_ − RRCS_inactive_, reflects the change of RRCS for a residue pair from inactive- to active- state (Figure 2b-d and Figure 2—figure supplement 1b). To identify residue pairs with conserved conformational rearrangements upon activation across class A GPCRs, two rounds of selections (Figure 2d and Figure 2—Source data 1) were performed: (i) identification of conserved rearrangements of residue contacts upon activation for six receptors (bRho, β_2_AR, M2R, μOR, A_2A_R and κ-OR), *i.e*., equivalent residue pairs show a similar and substantial change in RRCS between the active and inactive state structure of each of the six receptors (the same sign of ΔRRCS and |ΔRRCS| > 0.2 for all receptors) and (ii) family-wide RRCS comparison between the 142 inactive and 27 active state structures to identify residues pairs of statistically significant different (P<0.001; two sample *t*-test) RRCS upon activation.

#### Round 1

Identification of conserved rearrangements of residue contacts. Six receptors with available inactive- and active-state structures were analysed using ΔRRCS to identify residue pairs that share similar conformational changes. Twelve representative crystal structures (high-resolution, no mutation or one mutation in TMD without affecting receptor signalling) were chosen in this stage: 6 inactive state structures (PDB codes 1GZM for bRho, 2RH1 for β_2_AR, 3UON for M2R, 4DKL for μOR, 3EML for A_2A_R and 4DJH for κ-OR) and 6 active state structures (3PQR for bRho, 3SN6 for β_2_AR, 4MQS for M2R, 5C1M for μOR, 5G53 for A_2A_R and 6B73 for κ-OR) (Figure 2d, Figure 2—figure supplement 1c and Figure 2—Source data 1). Each receptor has approximately 600 residues pairs that have positive RRCS. Roughly one quarter are newly formed during receptor activation (RRCS_inactive_ =0 & RRCS_active_ >0); another quarter lose their contacts upon receptor activation (RRCS_inactive_ >0 & RRCS_active_ =0); and the remaining appear in both the inactive- or active- state structures (RRCS_inactive_ >0 & RRCS_active_ >0), the contact rearrangement of which can only be reflected by ΔRRCS, but not Boolean descriptors.

To identify residue pairs that share conserved rearrangements of residue contacts upon activation, two steps are performed to qualify residue pairs for the next round. Firstly, residue pairs with same sign of ΔRRCS and |ΔRRCS| > 0.2 for all six receptors were identified. There are 32 intra-receptor residues pairs (1×49:7×50, 1×53:7×53, 1×53:7×54, 2×37:2×40, 2×42:4×45, 2×43:7×53, 2×45:4×50, 2×46:2×50, 2×50:3×39, 2×57:7×42, 3×40:6×48, 3×43:6×40, 3×43:6×41, 3×43:7×49, 3×43:7×53, 3×46:6×37, 3×46:7×53, 3×49:3×50, 3×50:3×53, 3×50:6×37, 3×0:7×53, 3×51:5×57, 5×51:6×44, 5×58:6×40, 5×62:6×37, 6×40:7×49, 6×44:6×48, 7×50:7×55, 7×52:7×53, 7×53:8×50, 7×54:8×50 and 7×54:8×51) and 5 receptor-G protein/its mimetic residue pairs (3×50:G protein, 3×53:G protein, 3×54:G protein, 5×61:G protein and 6×33:G protein) that meet this criterion. Secondly, we also investigated residue pairs with ΔRRCS that are conserved in five receptors (*i.e.*, with one receptor as exception). Considering there is no Na^+^ pocket for rhodopsin, 3 residue pairs (2×50:7×49, 6×44:7×45, 6×48:7×45) around Na^+^ pocket were analysed for five receptors but not bRho. Additionally, 3 residue pairs have 0 (3×46:3×50, 5×55:6×41) or negative (7×45:7×49) ΔRRCS for κ-OR but positive ΔRRCS for the other five receptors. As for 3446:3450, nanobody-stabilized active structures (β_2_AR: 3P0G, 4LDO, 4LDL, 4LDE, 4QKX; and μOR: 5C1M) generally have lower contact scores (<0.4) compared with G protein-bound active-state structures (2.17 for 3SN6 of β_2_AR, 2.57 for 5G53 of A_2A_R and 6.93 for 3PQR of bRho). For these residue pairs, we added newly determined G_i_-bound active receptors A_1A_R and 5-HT_1B_ and found they have positive ΔRRCS, like other five receptors (Figure 4—figure supplements 1 and 2). Thus, these three residue pairs (3×46:3×50, 5×55:6×41 and 7×45:7×49) were retained. Totally, 6 residue pairs with conserved ΔRRCS in five receptors were rescued. Taken together, 38 intra-receptor residue pairs and 5 receptor-G protein/its mimetic residue pairs were identified to have conserved rearrangements of residue contacts upon activation.

#### Round 2

Family-wide conservation analysis of residue contact pattern. To investigate the conservation of residue contact pattern for the 38 intra-receptor residue pairs across these functionally diverse receptors, two-tailed unpaired *t*-test between inactive state (142 inactive structures from 38 receptors) and active state (27 active structures from 8 receptors) groups were performed (Figure 2d and Figure 2—Source data 2). Thirty one residue pairs have significantly different RRCS between inactive- and active-state (P<10^−5^). As rhodopsin lacks Na^+^ pocket, all rhodopsin structures were neglected in the analysis of 3 residue pairs around Na^+^ pocket (2×50:7×49, 6×44:7×45 and 6×48:7×45), which have good P value (<10^−3^) for these non-rhodopsin class A GPCRs. 4 residue pairs were filtered out in this round due to their poor P value, *i.e.*, there are no statistically significant difference in RRCS between inactive and active states (P=0.01 for 2×37:2×40, 0.96 for 2×42:4×45, 0.02 for 2×45:4×50 and 0.014 for 2×57:7×42).

Finally, 34 intra-receptor residue pairs (Figure 2d, Figure 4—figure supplements 1 and 2) and 5 receptor-G protein residue pairs were identified with conserved rearrangements of residue contacts upon activation, including all six residues pairs identified by the previous RC approaches ^8^.

### Sequence analysis of class A GPCRs

The alignment of 286 non-olfactory, class A human GPCRs were obtained from the GPCRdb^35, 62^. The distribution of sequence similarity/identity across class A GPCRs were extracted from the sequence similarity/identity matrix for different structural regions by using “Similarity matrix” tool in GPCRdb. The sequence conservation score (Figure 1—figure supplement 1) for all residue positions across 286 non-olfactory class A GPCRs were evaluated by the Protein Residue Conservation Prediction^56^ tool with scoring method “property entropy”^57^. Sequence conservation analysis (Figure 7—figure supplement 2) were visualized by WebLogo3^73^ with sequence alignment files from GPCRdb as the input.

### CAM/CIM in class A GPCRs

For the 14 hub residues in the universal activation pathway, we collected the functional mutation data from literature and GPCRdb^35, 62^. Mutations with “more than two fold-increase in basal activity/constitutively active” or “abolished effect” compared to the wild-type receptor were selected. Together, 272 mutations from 41 class A GPCRs on the 14 hub residues were collected, including the mutations we designed and validated in this work (Figure 7—source data 1).

### Disease-associated mutations in class A GPCRs

To reveal the relationship between disease-associated mutations and associated phenotypes of different transmembrane regions^74–77^, we collected disease-associated mutation information for all 286 non-olfactory class A GPCRs by database integration and literature investigation. Four commonly used databases (UniProt^58^, OMIM^59^, Ensembl^60^ and GPCRdb^54–55^) were first filtered by disease mutations and then merged. Totally 435 disease mutations from 61 class A GPCRs were collected (Figure 1—Source data 2).

### Pathway-guided CAM/CIM design in A_2A_R

We designed mutations for a prototypical receptor A_2A_R, guided by the universal activation pathway, aiming to get constitutively active/inactive receptor. Mutations that can either stabilize active or inactive state structures of A_2A_R or promote/block the conformational change upon activation were designed (Figure 6c and Figure 6—figure supplement 1) and tested by functional cAMP accumulation assays. The inactive state structure 3EML and active state structure 5G53 were used. *In silico* mutagenesis was performed by Residue Scanning module in BioLuminate^61^. Sidechain prediction with backbone sampling and a cut-off value of 6Å were applied during the scanning. ΔStability is the change of receptor stability when introducing a mutation. We filtered the mutations by one of the following criteria: (i) ΔStability in active and inactive structures have opposite sign; or (ii) ΔStability in active and inactive structures have the same sign, but favourable interactions such as hydrogen bonds, salt bridge or pi-pi stacking exist in only one structure that can promote/block the conformational change upon activation. Totally, 15 and 20 mutations were predicted to be CAMs and CIMs, respectively. (Figure 6c and Figure 6—figure supplement 1).

### cAMP accumulation assay

The desired mutations were introduced into amino-terminally Flago® tag-labeled human A_2A_R in the pcDNA3.1 vector (Invitrogen, Carlsbad, CA, USA). This construct displayed equivalent pharmacological features to that of untagged human receptor based on radioligand binding and cAMP assays^78^. The mutants were constructed by PCR-based site-directed mutagenesis (Muta-directTM kit, Beijing SBS Genetech Co., Ltd., China). Sequences of receptor clones were confirmed by DNA sequencing. HEK-293 cells were seeded onto 6-well cell culture plates. After overnight culture, the cells were transiently transfected with WT or mutant DNA using Lipofectamine 2000 transfection reagent (Invitrogen). After 24 h, the transfected cells were seeded onto 384-well plates (3,000 cells per well). cAMP accumulation was measured using the LANCE cAMP kit (PerkinElmer, Boston, MA, USA) according to the manufacturer’s instructions. Briefly, transfected cells were incubated for 40 min in assay buffer (DMEM, 1 mM 3-isobutyl-1-methylxanthine) with different concentrations of agonist [CGS21680 (179 pM to 50 μM)]. The reactions were stopped by addition of lysis buffer containing LANCE reagents. Plates were then incubated for 60 min at room temperature and time-resolved FRET signals were measured at 625 nm and 665 nm by an EnVision multilabel plate reader (PerkinElmer). The cAMP response is depicted relative to the maximal response of CGS21680 (100%) at the WT A_2A_R.

### CGS21680 binding assay

CGS21680 (a specific adenosine A_2A_ subtype receptor agonist) binding was analyzed using plasma membranes prepared from HEK-293 cells transiently expressing WT and mutant A_2A_Rs. Approximately 1.2 × 10^8^ transfected HEK-293 cells were harvested, suspended in 10 ml ice-cold membrane buffer (50 mM Tris-HCl, pH 7.4) and centrifuged for 5 min at 700 ***g***. The resulting pellet was resuspended in ice-cold membrane buffer, homogenized by Dounce Homogenizer (Wheaton, Millville, NJ, USA) and centrifuged for 20 min at 50,000 ***g***. The pellet was resuspended, homogenized, centrifuged again and the precipitate containing the plasma membranes was then suspended in the membrane buffer containing protease inhibitor (Sigma-Aldrich, St. Louis, MO, USA) and stored at −80°C. Protein concentration was determined using a protein BCA assay kit (Pierce Biotechnology, Pittsburgh, PA, USA). For homogeneous binding, cell membrane homogenates (10 μg protein per well) were incubated in membrane binding buffer (50 mM Tris-HCl, 10 mM NaCl, 0.1 mM EDTA, pH 7.4) with constant concentration of [^3^H]-CGS21680 (1 nM, PerkinElmer) and serial dilutions of unlabeled CGS21680 (0.26 nM to 100 μM) at room temperature for 3 h. Nonspecific binding was determined in the presence of 100 μM CGS21680. Following incubation, the samples were filtered rapidly in vacuum through glass fiber filter plates (PerkinElmer). After soaking and rinsing 4 times with ice-cold PBS, the filters were dried and counted for radioactivity in a MicroBeta2 scintillation counter (PerkinElmer).

### Surface expression of A_2A_Rs

HEK-293 cells were seeded into 6-well plate and incubated overnight. After transient transfection with WT or mutant plasmids for 24 h, the cells were collected and blocked with 5% BSA in PBS at room temperature for 15 min and incubated with primary anti-Flag antibody (1:100, Sigma-Aldrich) at room temperature for 1 h. The cells were then washed three times with PBS containing 1% BSA followed by 1 h incubation with anti-rabbit Alexa-488-conjugated secondary antibody (1:1000, Cell Signaling Technology, Danvers, MA, USA) at 4°C in the dark. After three washes, the cells were resuspended in 200 μl of PBS containing 1% BSA for detection in a NovoCyte flow cytometer (ACEA Biosciences, San Diego, CA, USA) utilizing laser excitation and emission wavelengths of 488 nm and 519 nm, respectively. For each assay point, approximately 15,000 cellular events were collected, and the total fluorescence intensity of positive expression cell population was calculated.

## Data and materials availability

The open source code is available at GitHub (https://github.com/zhaolabSHT/RRCS). For availability of codes that were developed in-house, please contacts the corresponding authors. All data is available in the main text or the source data.

## Acknowledgments

This work was partially supported by National Key R&D Program of China grants 2016YFC0905900 (S.Z.) and 2018YFA0507000 (S.Z. and M.-W.W.), National Mega R&D Program for Drug Discovery grants 2018ZX09711002-002-005 (D.H.Y.) and 2018ZX09735-001 (M.-W.W), National Natural Science Foundation of China grants 21704064 (Q.Z.), 81573479 (D.H.Y.) and 81773792 (D.H.Y.), Shanghai Science & Technology Development Fund grants 16ZR1448500 (S.Z.) and 16ZR1407100 (A.T.D.), Young Talent Program of Shanghai (S.Z.), Novo Nordisk-CAS Research Fund grant NNCAS-2017-1-CC (D.H.Y.), the Medical Research Council MC_U105185859 (M.M.B.), and annual overhead support from ShanghaiTech University and Chinese Academy of Sciences. We thank A. Sali, M.A. Hanson and A.J. Venkatakrishnan for valuable discussions, Y.M. Xu for technical assistance, and A.P. IJzerman for providing the WT A_2A_R plasmid.

## Authors contributions

Q.Z. and S.Z. conceived the project; M.-W.W. and M.M.B. expanded the scope of the project. Q.Z. performed computational studies. Q.Z., D.H.Y., S.Z. and M.M.B. designed the mutagenesis experiments. M.W. helped with data collection and initial mutagenesis study. Y.G. wrote the code for RRCS calculation. W.J.G., L.Z., X.Q.C., A.T.D. and D.H.Y. conducted mutagenesis and pharmacology experiments with the supervision of M.-W.W. E.S., Z.-J.L. and R.C.S. provided academic support or technical inputs to the project. Q.Z., D.H.Y., M.M.B., M-W.W. and S.Z. interpreted the data and wrote the manuscript. S.Z. and M.-W.W. managed the entire project.

## References

1. Deupi, X.; Standfuss, J., Structural insights into agonist-induced activation of G-protein-coupled receptors. Curr Opin Struct Biol 2011, 21 (4), 541–51. https://doi.org/10.1016/j.sbi.2011.06.002.

2. Venkatakrishnan, A. J.; Deupi, X.; Lebon, G.; Tate, C. G.; Schertler, G. F.; Babu, M. M., Molecular signatures of G-protein-coupled receptors. Nature 2013, 494 (7436), 185–94. https://doi.org/10.1038/nature11896.

3. Katritch, V.; Cherezov, V.; Stevens, R. C., Structure-function of the G protein-coupled receptor superfamily. Annu Rev Pharmacol Toxicol 2013, 53, 531–56. https://doi.org/10.1146/annurev-pharmtox-032112-135923.

4. Erlandson, S. C.; McMahon, C.; Kruse, A. C., Structural Basis for G Protein-Coupled Receptor Signaling. Annu Rev Biophys 2018. https://doi.org/10.1146/annurev-biophys-070317-032931.

5. Weis, W. I.; Kobilka, B. K., The Molecular Basis of G Protein-Coupled Receptor Activation. Annual review of biochemistry 2018, 87, 897–919. https://doi.org/10.1146/annurev-biochem-060614-033910.

6. Ghosh, E.; Kumari, P.; Jaiman, D.; Shukla, A. K., Methodological advances: the unsung heroes of the GPCR structural revolution. Nat Rev Mol Cell Biol 2015, 16 (2), 69–81. https://doi.org/10.1038/nrm3933.

7. Cooke, R. M.; Brown, A. J.; Marshall, F. H.; Mason, J. S., Structures of G protein-coupled receptors reveal new opportunities for drug discovery. Drug Discov Today 2015, 20 (11), 1355–64. https://doi.org/10.1016/j.drudis.2015.08.003.

8. Venkatakrishnan, A. J.; Deupi, X.; Lebon, G.; Heydenreich, F. M.; Flock, T.; Miljus, T.; Balaji, S.; Bouvier, M.; Veprintsev, D. B.; Tate, C. G.; Schertler, G. F.; Babu, M. M., Diverse activation pathways in class A GPCRs converge near the G-protein-coupling region. Nature 2016, 536 (7617), 484–7. https://doi.org/10.1038/nature19107.

9. Wootten, D.; Christopoulos, A.; Marti-Solano, M.; Babu, M. M.; Sexton, P. M., Mechanisms of signalling and biased agonism in G protein-coupled receptors. Nat Rev Mol Cell Biol 2018, 19 (10), 638–653. https://doi.org/10.1038/s41580-018-0049-3.

10. Flock, T.; Hauser, A. S.; Lund, N.; Gloriam, D. E.; Balaji, S.; Babu, M. M., Selectivity determinants of GPCR-G-protein binding. Nature 2017, 545 (7654), 317–322. https://doi.org/10.1038/nature22070.

11. Hofmann, K. P.; Scheerer, P.; Hildebrand, P. W.; Choe, H. W.; Park, J. H.; Heck, M.; Ernst, O. P., A G protein-coupled receptor at work: the rhodopsin model. Trends Biochem Sci 2009, 34 (11), 540–52. https://doi.org/10.1016/j.tibs.2009.07.005.

12. Katritch, V.; Cherezov, V.; Stevens, R. C., Diversity and modularity of G protein-coupled receptor structures. Trends Pharmacol Sci 2012, 33 (1), 17–27. https://doi.org/10.1016/j.tips.2011.09.003.

13. Dror, R. O.; Green, H. F.; Valant, C.; Borhani, D. W.; Valcourt, J. R.; Pan, A. C.; Arlow, D. H.; Canals, M.; Lane, J. R.; Rahmani, R.; Baell, J. B.; Sexton, P. M.; Christopoulos, A.; Shaw, D. E., Structural basis for modulation of a G-protein-coupled receptor by allosteric drugs. Nature 2013, 503 (7475), 295–9. https://doi.org/10.1038/nature12595.

14. Ye, L.; Van Eps, N.; Zimmer, M.; Ernst, O. P.; Prosser, R. S., Activation of the A2A adenosine G-protein-coupled receptor by conformational selection. Nature 2016, 533 (7602), 265–8. https://doi.org/10.1038/nature17668.

15. Schonegge, A. M.; Gallion, J.; Picard, L. P.; Wilkins, A. D.; Le Gouill, C.; Audet, M.; Stallaert, W.; Lohse, M. J.; Kimmel, M.; Lichtarge, O.; Bouvier, M., Evolutionary action and structural basis of the allosteric switch controlling beta2AR functional selectivity. Nat Commun 2017, 8 (1), 2169. https://doi.org/10.1038/s41467-017-02257-x.

16. Latorraca, N. R.; Venkatakrishnan, A. J.; Dror, R. O., GPCR Dynamics: Structures in Motion. Chem Rev 2017, 117 (1), 139–155. https://doi.org/10.1021/acs.chemrev.6b00177.

17. Thal, D. M.; Glukhova, A.; Sexton, P. M.; Christopoulos, A., Structural insights into G-protein-coupled receptor allostery. Nature 2018, 559 (7712), 45–53. https://doi.org/10.1038/s41586-018-0259-z.

18. Venkatakrishnan, A. J.; Ma, A. K.; Fonseca, R.; Latorraca, N. R.; Kelly, B.; Betz, R. M.; Asawa, C.; Kobilka, B. K.; Dror, R. O., Diverse GPCRs exhibit conserved water networks for stabilization and activation. Proc Natl Acad Sci U S A 2019, 116 (8), 3288–3293. https://doi.org/10.1073/pnas.1809251116.

19. Yuan, S.; Vogel, H.; Filipek, S., The role of water and sodium ions in the activation of the mu-opioid receptor. Angew Chem Int Ed Engl 2013, 52 (38), 10112–5. https://doi.org/10.1002/anie.201302244.

20. Nygaard, R.; Frimurer, T. M.; Holst, B.; Rosenkilde, M. M.; Schwartz, T. W., Ligand binding and micro-switches in 7TM receptor structures. Trends Pharmacol Sci 2009, 30 (5), 249–59. https://doi.org/10.1016/j.tips.2009.02.006.

21. Trzaskowski, B.; Latek, D.; Yuan, S.; Ghoshdastider, U.; Debinski, A.; Filipek, S., Action of molecular switches in GPCRs--theoretical and experimental studies. Curr Med Chem 2012, 19 (8), 1090–109.

22. Tehan, B. G.; Bortolato, A.; Blaney, F. E.; Weir, M. P.; Mason, J. S., Unifying family A GPCR theories of activation. Pharmacol Ther 2014, 143 (1), 51–60. https://doi.org/10.1016/j.pharmthera.2014.02.004.

23. Rasmussen, S. G.; DeVree, B. T.; Zou, Y.; Kruse, A. C.; Chung, K. Y.; Kobilka, T. S.; Thian, F. S.; Chae, P. S.; Pardon, E.; Calinski, D.; Mathiesen, J. M.; Shah, S. T.; Lyons, J. A.; Caffrey, M.; Gellman, S. H.; Steyaert, J.; Skiniotis, G.; Weis, W. I.; Sunahara, R. K.; Kobilka, B. K., Crystal structure of the beta2 adrenergic receptor-Gs protein complex. Nature 2011, 477 (7366), 549–55. https://doi.org/10.1038/nature10361.

24. Katritch, V.; Fenalti, G.; Abola, E. E.; Roth, B. L.; Cherezov, V.; Stevens, R. C., Allosteric sodium in class A GPCR signaling. Trends Biochem Sci 2014, 39 (5), 233–44. https://doi.org/10.1016/j.tibs.2014.03.002.

25. Bhattacharya, S.; Vaidehi, N., Differences in allosteric communication pipelines in the inactive and active states of a GPCR. Biophys J 2014, 107 (2), 422–434. https://doi.org/10.1016/j.bpj.2014.06.015.

26. Rodriguez, G. J.; Yao, R.; Lichtarge, O.; Wensel, T. G., Evolution-guided discovery and recoding of allosteric pathway specificity determinants in psychoactive bioamine receptors. Proc Natl Acad Sci U S A 2010, 107 (17), 7787–92. https://doi.org/10.1073/pnas.0914877107.

27. Manglik, A.; Kim, T. H.; Masureel, M.; Altenbach, C.; Yang, Z.; Hilger, D.; Lerch, M. T.; Kobilka, T. S.; Thian, F. S.; Hubbell, W. L.; Prosser, R. S.; Kobilka, B. K., Structural Insights into the Dynamic Process of beta2-Adrenergic Receptor Signaling. Cell 2015, 161 (5), 1101–1111. https://doi.org/10.1016/j.cell.2015.04.043.

28. Wescott, M. P.; Kufareva, I.; Paes, C.; Goodman, J. R.; Thaker, Y.; Puffer, B. A.; Berdougo, E.; Rucker, J. B.; Handel, T. M.; Doranz, B. J., Signal transmission through the CXC chemokine receptor 4 (CXCR4) transmembrane helices. Proc. Natl. Acad. Sci. U. S. A. 2016, 113 (35), 9928–33. https://doi.org/10.1073/pnas.1601278113.

29. Furness, S. G. B.; Liang, Y. L.; Nowell, C. J.; Halls, M. L.; Wookey, P. J.; Dal Maso, E.; Inoue, A.; Christopoulos, A.; Wootten, D.; Sexton, P. M., Ligand-Dependent Modulation of G Protein Conformation Alters Drug Efficacy. Cell 2016, 167 (3), 739–749e11. https://doi.org/10.1016/j.cell.2016.09.021.

30. Sung, Y. M.; Wilkins, A. D.; Rodriguez, G. J.; Wensel, T. G.; Lichtarge, O., Intramolecular allosteric communication in dopamine D2 receptor revealed by evolutionary amino acid covariation. Proc Natl Acad Sci U S A 2016, 113 (13), 3539–44. https://doi.org/10.1073/pnas.1516579113.

31. Vaidehi, N.; Bhattacharya, S., Allosteric communication pipelines in G-protein-coupled receptors. Current opinion in pharmacology 2016, 30, 76–83. https://doi.org/10.1016/j.coph.2016.07.010.

32. Che, T.; Majumdar, S.; Zaidi, S. A.; Ondachi, P.; McCorvy, J. D.; Wang, S.; Mosier, P. D.; Uprety, R.; Vardy, E.; Krumm, B. E.; Han, G. W.; Lee, M. Y.; Pardon, E.; Steyaert, J.; Huang, X. P.; Strachan, R. T.; Tribo, A. R.; Pasternak, G. W.; Carroll, F. I.; Stevens, R. C.; Cherezov, V.; Katritch, V.; Wacker, D.; Roth, B. L., Structure of the Nanobody-Stabilized Active State of the Kappa Opioid Receptor. Cell 2018, 172 (1-2), 55–67 e15. https://doi.org/10.1016/j.cell.2017.12.011.

33. Eddy, M. T.; Lee, M. Y.; Gao, Z. G.; White, K. L.; Didenko, T.; Horst, R.; Audet, M.; Stanczak, P.; McClary, K. M.; Han, G. W.; Jacobson, K. A.; Stevens, R. C.; Wuthrich, K., Allosteric Coupling of Drug Binding and Intracellular Signaling in the A2A Adenosine Receptor. Cell 2018, 172 (1-2), 68–80 e12. https://doi.org/10.1016/j.cell.2017.12.004.

34. Ngo, T.; Ilatovskiy, A. V.; Stewart, A. G.; Coleman, J. L.; McRobb, F. M.; Riek, R. P.; Graham, R. M.; Abagyan, R.; Kufareva, I.; Smith, N. J., Orphan receptor ligand discovery by pickpocketing pharmacological neighbors. Nat Chem Biol 2017, 13 (2), 235–242. https://doi.org/10.1038/nchembio.2266.

35. Isberg, V.; Mordalski, S.; Munk, C.; Rataj, K.; Harpsoe, K.; Hauser, A. S.; Vroling, B.; Bojarski, A. J.; Vriend, G.; Gloriam, D. E., GPCRdb: an information system for G protein-coupled receptors. Nucleic Acids Res 2016, 44 (D1), D356–64. https://doi.org/10.1093/nar/gkv1178.

36. Valentin-Hansen, L.; Holst, B.; Frimurer, T. M.; Schwartz, T. W., PheVI:09 (Phe6.44) as a sliding microswitch in seven-transmembrane (7TM) G protein-coupled receptor activation. J Biol Chem 2012, 287 (52), 43516–26. https://doi.org/10.1074/jbc.M112.395137.

37. Rovati, G. E.; Capra, V.; Neubig, R. R., The highly conserved DRY motif of class A G protein-coupled receptors: beyond the ground state. Mol Pharmacol 2007, 71 (4), 959–64. https://doi.org/10.1124/mol.106.029470.

38. Glukhova, A.; Draper-Joyce, C. J.; Sunahara, R. K.; Christopoulos, A.; Wootten, D.; Sexton, P. M., Rules of engagement: GPCRs and G proteins. ACS Pharmacology & Translational Science 2018. https://doi.org/10.1021/acsptsci.8b00026.

39. Hulme, E. C., GPCR activation: a mutagenic spotlight on crystal structures. Trends Pharmacol Sci 2013, 34 (1), 67–84. https://doi.org/10.1016/j.tips.2012.11.002.

40. Erdelyi, L. S.; Mann, W. A.; Morris-Rosendahl, D. J.; Gross, U.; Nagel, M.; Varnai, P.; Balla, A.; Hunyady, L., Mutation in the V2 vasopressin receptor gene, AVPR2, causes nephrogenic syndrome of inappropriate diuresis. Kidney Int 2015, 88 (5), 1070–8. https://doi.org/10.1038/ki.2015.181.

41. Pasel, K.; Schulz, A.; Timmermann, K.; Linnemann, K.; Hoeltzenbein, M.; Jaaskelainen, J.; Gruters, A.; Filler, G.; Schoneberg, T., Functional characterization of the molecular defects causing nephrogenic diabetes insipidus in eight families. J Clin Endocrinol Metab 2000, 85 (4), 1703–10. https://doi.org/10.1210/jcem.85.4.6507.

42. Napier, M. L.; Durga, D.; Wolsley, C. J.; Chamney, S.; Alexander, S.; Brennan, R.; Simpson, D. A.; Silvestri, G.; Willoughby, C. E., Mutational Analysis of the Rhodopsin Gene in Sector Retinitis Pigmentosa. Ophthalmic Genet 2015, 36 (3), 239–43. https://doi.org/10.3109/13816810.2014.958862.

43. Wingler, L. M.; Elgeti, M.; Hilger, D.; Latorraca, N. R.; Lerch, M. T.; Staus, D. P.; Dror, R. O.; Kobilka, B. K.; Hubbell, W. L.; Lefkowitz, R. J., Angiotensin Analogs with Divergent Bias Stabilize Distinct Receptor Conformations. Cell 2019, 176 (3), 468–+. https://doi.org/10.1016/j.cell.2018.12.005.

44. Tan, L.; Yan, W.; McCorvy, J. D.; Cheng, J., Biased Ligands of G Protein-Coupled Receptors (GPCRs): Structure-Functional Selectivity Relationships (SFSRs) and Therapeutic Potential. J Med Chem 2018. https://doi.org/10.1021/acs.jmedchem.8b00435.

45. Smith, J. S.; Lefkowitz, R. J.; Rajagopal, S., Biased signalling: from simple switches to allosteric microprocessors. Nat Rev Drug Discov 2018. https://doi.org/10.1038/nrd.2017.229.

46. Schmid, C. L.; Kennedy, N. M.; Ross, N. C.; Lovell, K. M.; Yue, Z. Z.; Morgenweck, J.; Cameron, M. D.; Bannister, T. D.; Bohn, L. M., Bias Factor and Therapeutic Window Correlate to Predict Safer Opioid Analgesics. Cell 2017, 171 (5), 1165–+. https://doi.org/10.1016/j.cell.2017.10.035.

47. Wootten, D.; Reynolds, C. A.; Smith, K. J.; Mobarec, J. C.; Koole, C.; Savage, E. E.; Pabreja, K.; Simms, J.; Sridhar, R.; Furness, S. G.; Liu, M.; Thompson, P. E.; Miller, L. J.; Christopoulos, A.; Sexton, P. M., The Extracellular Surface of the GLP-1 Receptor Is a Molecular Trigger for Biased Agonism. Cell 2016, 165 (7), 1632–43. https://doi.org/10.1016/j.cell.2016.05.023.

48. Lu, S.; Zhang, J., Small Molecule Allosteric Modulators of G-Protein-Coupled Receptors: Drug-Target Interactions. J Med Chem 2018. https://doi.org/10.1021/acs.jmedchem.7b01844.

49. Gregorio, G. G.; Masureel, M.; Hilger, D.; Terry, D. S.; Juette, M.; Zhao, H.; Zhou, Z.; Perez-Aguilar, J. M.; Hauge, M.; Mathiasen, S.; Javitch, J. A.; Weinstein, H.; Kobilka, B. K.; Blanchard, S. C., Single-molecule analysis of ligand efficacy in beta2AR-G-protein activation. Nature 2017, 547 (7661), 68–73. https://doi.org/10.1038/nature22354.

50. Isogai, S.; Deupi, X.; Opitz, C.; Heydenreich, F. M.; Tsai, C. J.; Brueckner, F.; Schertler, G. F.; Veprintsev, D. B.; Grzesiek, S., Backbone NMR reveals allosteric signal transduction networks in the beta1-adrenergic receptor. Nature 2016, 530 (7589), 237–41. https://doi.org/10.1038/nature16577.

51. Lee, M. H.; Appleton, K. M.; Strungs, E. G.; Kwon, J. Y.; Morinelli, T. A.; Peterson, Y. K.; Laporte, S. A.; Luttrell, L. M., The conformational signature of beta-arrestin2 predicts its trafficking and signalling functions. Nature 2016, 531 (7596), 665–8. https://doi.org/10.1038/nature17154.

52. Tian, H.; Furstenberg, A.; Huber, T., Labeling and Single-Molecule Methods To Monitor G Protein-Coupled Receptor Dynamics. Chem Rev 2017, 117 (1), 186–245. https://doi.org/10.1021/acs.chemrev.6b00084.

53. Dror, R. O.; Pan, A. C.; Arlow, D. H.; Borhani, D. W.; Maragakis, P.; Shan, Y.; Xu, H.; Shaw, D. E., Pathway and mechanism of drug binding to G-protein-coupled receptors. Proc Natl Acad Sci U S A 2011, 108 (32), 13118–23. https://doi.org/10.1073/pnas.1104614108.

54. Hauser, A. S.; Chavali, S.; Masuho, I.; Jahn, L. J.; Martemyanov, K. A.; Gloriam, D. E.; Babu, M. M., Pharmacogenomics of GPCR Drug Targets. Cell 2018, 172 (1-2), 41–54 e19. https://doi.org/10.1016/j.cell.2017.11.033.

55. Pandy-Szekeres, G.; Munk, C.; Tsonkov, T. M.; Mordalski, S.; Harpsoe, K.; Hauser, A. S.; Bojarski, A. J.; Gloriam, D. E., GPCRdb in 2018: adding GPCR structure models and ligands. Nucleic Acids Res 2018, 46 (D1), D440–D446. https://doi.org/10.1093/nar/gkx1109.

56. Capra, J. A.; Singh, M., Predicting functionally important residues from sequence conservation. Bioinformatics 2007, 23 (15), 1875–82. https://doi.org/10.1093/bioinformatics/btm270.

57. Mirny, L. A.; Shakhnovich, E. I., Universally conserved positions in protein folds: reading evolutionary signals about stability, folding kinetics and function. J Mol Biol 1999, 291 (1), 177–96. https://doi.org/10.1006/jmbi.1999.2911.

58. The UniProt, C., UniProt: the universal protein knowledgebase. Nucleic Acids Res 2017, 45 (D1), D158–D169. https://doi.org/10.1093/nar/gkw1099.

59. Amberger, J.; Bocchini, C.; Hamosh, A., A new face and new challenges for Online Mendelian Inheritance in Man (OMIM(R)). Human mutation 2011, 32 (5), 564–7. https://doi.org/10.1002/humu.21466.

60. Zerbino, D. R.; Achuthan, P.; Akanni, W.; Amode, M. R.; Barrell, D.; Bhai, J.; Billis, K.; Cummins, C.; Gall, A.; Giron, C. G.; Gil, L.; Gordon, L.; Haggerty, L.; Haskell, E.; Hourlier, T.; Izuogu, O. G.; Janacek, S. H.; Juettemann, T.; To, J. K.; Laird, M. R.; Lavidas, I.; Liu, Z.; Loveland, J. E.; Maurel, T.; McLaren, W.; Moore, B.; Mudge, J.; Murphy, D. N.; Newman, V.; Nuhn, M.; Ogeh, D.; Ong, C. K.; Parker, A.; Patricio, M.; Riat, H. S.; Schuilenburg, H.; Sheppard, D.; Sparrow, H.; Taylor, K.; Thormann, A.; Vullo, A.; Walts, B.; Zadissa, A.; Frankish, A.; Hunt, S. E.; Kostadima, M.; Langridge, N.; Martin, F. J.; Muffato, M.; Perry, E.; Ruffier, M.; Staines, D. M.; Trevanion, S. J.; Aken, B. L.; Cunningham, F.; Yates, A.; Flicek, P., Ensembl 2018. Nucleic Acids Res 2018, 46 (D1), D754–D761. https://doi.org/10.1093/nar/gkx1098.

61. Beard, H.; Cholleti, A.; Pearlman, D.; Sherman, W.; Loving, K. A., Applying physics-based scoring to calculate free energies of binding for single amino acid mutations in protein-protein complexes. PLoS One 2013, 8 (12), e82849. https://doi.org/10.1371/journal.pone.0082849.

62. Isberg, V.; de Graaf, C.; Bortolato, A.; Cherezov, V.; Katritch, V.; Marshall, F. H.; Mordalski, S.; Pin, J. P.; Stevens, R. C.; Vriend, G.; Gloriam, D. E., Generic GPCR residue numbers - aligning topology maps while minding the gaps. Trends Pharmacol Sci 2015, 36 (1), 22–31. https://doi.org/10.1016/j.tips.2014.11.001.

63. Ballesteros, J. A.; Weinstein, H., Integrated methods for the construction of three-dimensional models and computational probing of structure-function relations in G protein coupled receptors. In Receptor Molecular Biology, Sealfon, S. C., Ed. Academic Press: 1995; Vol. 25, pp 366–428.

64. Eldridge, M. D.; Murray, C. W.; Auton, T. R.; Paolini, G. V.; Mee, R. P., Empirical scoring functions: I. The development of a fast empirical scoring function to estimate the binding affinity of ligands in receptor complexes. Journal of computer-aided molecular design 1997, 11 (5), 425–45.

65. Verdonk, M. L.; Cole, J. C.; Hartshorn, M. J.; Murray, C. W.; Taylor, R. D., Improved protein-ligand docking using GOLD. Proteins 2003, 52 (4), 609–23. https://doi.org/10.1002/prot.10465.

66. Wang, S.; Sun, S.; Li, Z.; Zhang, R.; Xu, J., Accurate De Novo Prediction of Protein Contact Map by Ultra-Deep Learning Model. PLoS Comput Biol 2017, 13 (1), e1005324. https://doi.org/10.1371/journal.pcbi.1005324.

67. Kufareva, I.; Abagyan, R., Methods of Protein Structure Comparison. In Homology Modeling: Methods and Protocols, Orry, A. J. W.; Abagyan, R., Eds. Humana Press: Totowa, NJ, 2012; pp 231–257.

68. Kufareva, I.; Rueda, M.; Katritch, V.; Stevens, R. C.; Abagyan, R.; participants, G. D., Status of GPCR modeling and docking as reflected by community-wide GPCR Dock 2010 assessment. Structure 2011, 19 (8), 1108–26. https://doi.org/10.1016/j.str.2011.05.012.

69. Marsden, B.; Abagyan, R., SAD--a normalized structural alignment database: improving sequence-structure alignments. Bioinformatics 2004, 20 (15), 2333–44. https://doi.org/10.1093/bioinformatics/bth244.

70. Jones, T. A.; Zou, J. Y.; Cowan, S. W.; Kjeldgaard, M., Improved methods for building protein models in electron density maps and the location of errors in these models. Acta Crystallogr A 1991, 47 (Pt 2), 110–9. https://doi.org/10.1107/s0108767390010224.

71. Smart, O. S.; Horsky, V.; Gore, S.; Svobodova Varekova, R.; Bendova, V.; Kleywegt, G. J.; Velankar, S., Validation of ligands in macromolecular structures determined by X-ray crystallography. Acta Crystallogr D Struct Biol 2018, 74 (Pt 3), 228–236. https://doi.org/10.1107/S2059798318002541.

72. Kleywegt, G. J.; Harris, M. R.; Zou, J. Y.; Taylor, T. C.; Wahlby, A.; Jones, T. A., The Uppsala Electron-Density Server. Acta Crystallogr D Biol Crystallogr 2004, 60 (Pt 12 Pt 1), 2240–9. https://doi.org/10.1107/S0907444904013253.

73. Crooks, G. E.; Hon, G.; Chandonia, J. M.; Brenner, S. E., WebLogo: a sequence logo generator. Genome research 2004, 14 (6), 1188–90. https://doi.org/10.1101/gr.849004.

74. Vassart, G.; Costagliola, S., G protein-coupled receptors: mutations and endocrine diseases. Nat Rev Endocrinol 2011, 7 (6), 362–72. https://doi.org/10.1038/nrendo.2011.20.

75. Thompson, M. D.; Hendy, G. N.; Percy, M. E.; Bichet, D. G.; Cole, D. E., G protein-coupled receptor mutations and human genetic disease. Methods Mol Biol 2014, 1175, 153–87. https://doi.org/10.1007/978-1-4939-0956-8_8.

76. Tao, Y. X., Inactivating mutations of G protein-coupled receptors and diseases: structure-function insights and therapeutic implications. Pharmacol Ther 2006, 111 (3), 949–73. https://doi.org/10.1016/j.pharmthera.2006.02.008.

77. Tao, Y. X., Constitutive activation of G protein-coupled receptors and diseases: insights into mechanisms of activation and therapeutics. Pharmacol Ther 2008, 120 (2), 129–48. https://doi.org/10.1016/j.pharmthera.2008.07.005.

78. Massink, A.; Gutierrez-de-Teran, H.; Lenselink, E. B.; Ortiz Zacarias, N. V.; Xia, L.; Heitman, L. H.; Katritch, V.; Stevens, R. C.; AP, I. J., Sodium ion binding pocket mutations and adenosine A2A receptor function. Mol Pharmacol 2015, 87 (2), 305–13. https://doi.org/10.1124/mol.114.095737.

